# Fibrillarin-mediated ribosomal RNA maturation is a novel therapeutic vulnerability in triple-negative breast cancer

**DOI:** 10.1101/2025.02.23.639238

**Authors:** Camille Jouines, Piero Lo Monaco, Angéline Gaucherot, Julie Radermecker, Caroline Isaac, Fleur Bourdelais, Marie-Ambre Monet, Marion Meyer, Mounira Chalabi-Dchar, Flora Nguyen Van Long, Laury Baillon, Carine Froment, Julien Marcoux, Christophe Vanbelle, Tanguy Fenouil, Sébastien Durand, Stéphane Giraud, Jean-Jacques Diaz, Virginie Marcel, Frédéric Catez

## Abstract

Triple-negative breast cancer (TNBC) remains one of the most challenging breast cancer subtypes to treat due to the lack of effective therapeutic options. Ribosome biogenesis has recently emerged as a promising therapeutic target across various cancers. Despite the current targeting of ribosome biogenesis through RNA polymerase I (RNA Pol I) inhibition, we speculated that other factors essential for ribosome assembly, such as rRNA maturation factors, may also represent therapeutic targets in TNBC. Here, we demonstrate that ribosome biogenesis-related genes are notably overexpressed in TNBC compared to other breast cancer subtypes, highlighting its critical role in TNBC progression. Accordingly, we show that RNA Pol I inhibition exerts potent anti-proliferative effects in pre-clinical models of TNBC, both *in vitro* and *in vivo*. However, the DNA-damaging activity of RNA Pol I inhibitors raises safety concerns, highlighting the need for alternative strategies to inhibit ribosome biogenesis. To this end, we show that targeting a downstream rRNA maturation step, specifically pre-rRNA cleavage, by inhibiting the maturation factor Fibrillarin, also inhibits tumor growth in TNBC models. Notably, ribosome biogenesis inhibition, through either RNA Pol I or Fibrillarin targeting, induces cell cycle arrest without triggering significant cell death. These findings establish ribosome biogenesis as a therapeutic vulnerability in TNBC and identify rRNA maturation, and Fibrillarin in particular, as novel targets for potential therapeutic intervention.

**Significance:** Targeting ribosome biogenesis, through inhibition of either rRNA synthesis or maturation, induces anti-tumoral effects in TNBC, representing a novel therapeutic vulnerability with potential to improve patient outcomes.

## Introduction

Triple-negative breast cancer (TNBC), characterized by the absence of oestrogen and progesterone receptor expression, and of *HER2* gene amplification, accounts for 15-20% of breast cancer cases and represents a challenge in patient care. In particular, hormone therapies or HER2-targeted therapies are ineffective against TNBC, for which the standard-of-care remains neoadjuvant/preoperative chemotherapy followed by surgery and post-surgical treatments (1). The efficacy of these treatments is limited, and relapses are common due to the metastatic nature of TNBC. Patients harbouring TNBC have a 5-year survival rate ranging from 20% to 50%, depending on the metastatic site, highlighting the inter-tumor heterogeneity of TNBC (2). New therapies are available or under clinical evaluation, such as immunotherapy, PARP inhibitors, and the antibody-drug conjugate sacituzumab govitecan. However, response rates to these therapies remain below 30%, and adverse effects have been described (1,3).

New cancer therapeutic prospects have recently emerged from targeting ribosome biogenesis. Indeed, over activation of ribosome biogenesis is a hallmark of cancer cells, that is typically driven by the dysregulation of signalling pathways that control RNA Pol I activity, such as mTOR and c-MYC, p53 and PTEN (4). Importantly, tumor cells rely on the overactivation of ribosome biogenesis, which leads to an increase in the number of ribosomes, thus accelerating protein synthesis and metabolism, and favouring cell plasticity and adaptation to various stresses and changes in their environment, including chemotherapy (5). Ribosome biogenesis inhibition triggers ribosomal stress, which leads to a cytotoxic response that may include apoptosis, cell cycle arrest and autophagy depending on the cell type (6). Recently developed RNA Pol I inhibitors CX-5461 and BMH-21 provided the proof-of-concept that inhibiting ribosome biogenesis reduces cancer cell activity (7,8). CX-5461 and BMH-21 inhibits RNA Pol I by binding to G-quadruplex and GC-rich DNA sequences respectively, which are both abundant within the rDNA loci (7,8). However, off-target effects, specifically DNA damage, have been reported for CX-5461, likely due to DNA replication defects (9). Their therapeutic potential has so far been explored in various solid and haematological cancers, including leukaemia, lymphomas, myelomas, osteosarcoma, prostate cancer, ovarian cancer, glioma and neuroblastoma, but not in triple negative breast cancer (7,8,10–12). This dependency on ribosome biogenesis and its factors thus represents a vulnerability for cancer cells, which could be exploited as a therapeutic approach (5,13).

Though targeting RNA Pol I activity is the main strategy used to inhibit ribosome biogenesis, other factors could constitute promising candidates. Indeed, human ribosomes are assembled through a highly complex sequential and regulated process, that includes pre-rRNA endonuclease and exonuclease cleavages to release the mature 18S, 5.8S and 28S rRNAs, chemical modifications, folding and assembly with the 80 ribosomal proteins. These processes rely on hundreds of chaperones, nucleases, RNA helicases and non-coding C/D box and H/ACA small nucleolar RNAs (snoRNAs). Importantly, many of these ribosome biogenesis factors are essential to ribosome maturation, and may therefore be considered as therapeutic targets (13,14). For instance, inhibiting the non-coding U3 and U8 C/D box snoRNAs prevents the processing of pre-ribosomal RNAs into mature 18S and 28S rRNAs, and has anti-tumoral effects both *in vitro* and *in vivo* (*15*). Similarly, the Fibrillarin (FBL) protein, a key component of the C/D box snoRNP complexes, is also required for proper ribosome production in yeast and human cells (14,16–19). FBL contributes to mammary tumorigenesis and is overexpressed in several cancers, including breast cancer (17,20), acute myeloid leukaemia (21), prostate tumors (22) and liver cancer (23). In addition, in breast cancer, high FBL expression levels are associated with a poor prognosis (20).

Here, using pre-clinical *in vitro* and *in vivo* models, we demonstrate that ribosome biogenesis is a therapeutic vulnerability in TNBC, by inhibiting RNA Pol I activity and FBL-mediated rRNA maturation steps. We conclude that targeting rRNA maturation may offer a safer therapeutic strategy by avoiding the use of DNA binding compounds.

## Results

### The level of ribosome biogenesis-associated genes is higher in TNBC compared to other breast cancer subtypes

The molecular characterization of breast cancer subtypes has been instrumental in understanding their biology and identifying vulnerabilities for therapeutic intervention. As for other cancers, increased ribosome biogenesis is a hallmark of breast cancer cells (17,20,24), albeit it is unclear whether different levels of ribosome biogenesis are associated with specific breast cancer subtypes. To address this, we studied the level of expression of ribosome biogenesis-associated genes, used as a surrogate of ribosome biogenesis overactivation, as previously reported (20). We compared their levels across the different breast cancer subtypes using the BRCA gene expression dataset from The Cancer Genome Atlas (TCGA) public database (25). The ribosome biogenesis-associated gene signature encompassed 240 genes that defined the KEGG pathway “Ribosome Biogenesis in Eukaryotes” (Supplementary Table S1). Tumor samples (n = 739) were first categorized into four breast cancer subtypes based on the PAM50 molecular classification (Supplementary Table S2 and S3). The expression levels of ribosome biogenesis-associated genes were significantly higher in TNBC than in all other molecular breast cancer subtypes, including luminal A, luminal B and HER2-enriched (ANOVA, p = 2.79 x 10^−53^; Figure 1A). This finding was confirmed in a second group of tumor samples (n = 712) categorized according to their histological classification (*i.e.*, intrinsic breast cancer subtype) (ANOVA, p = 3.82 x 10^−29^; Figure 1B). Of note, despite using distinct parameters (histological and molecular) the groups of the two classifications overlapped (Supplementary Table S3). These results suggest that ribosome biogenesis is more active in TNBC than in other breast cancer subtypes.

**Figure 1.**
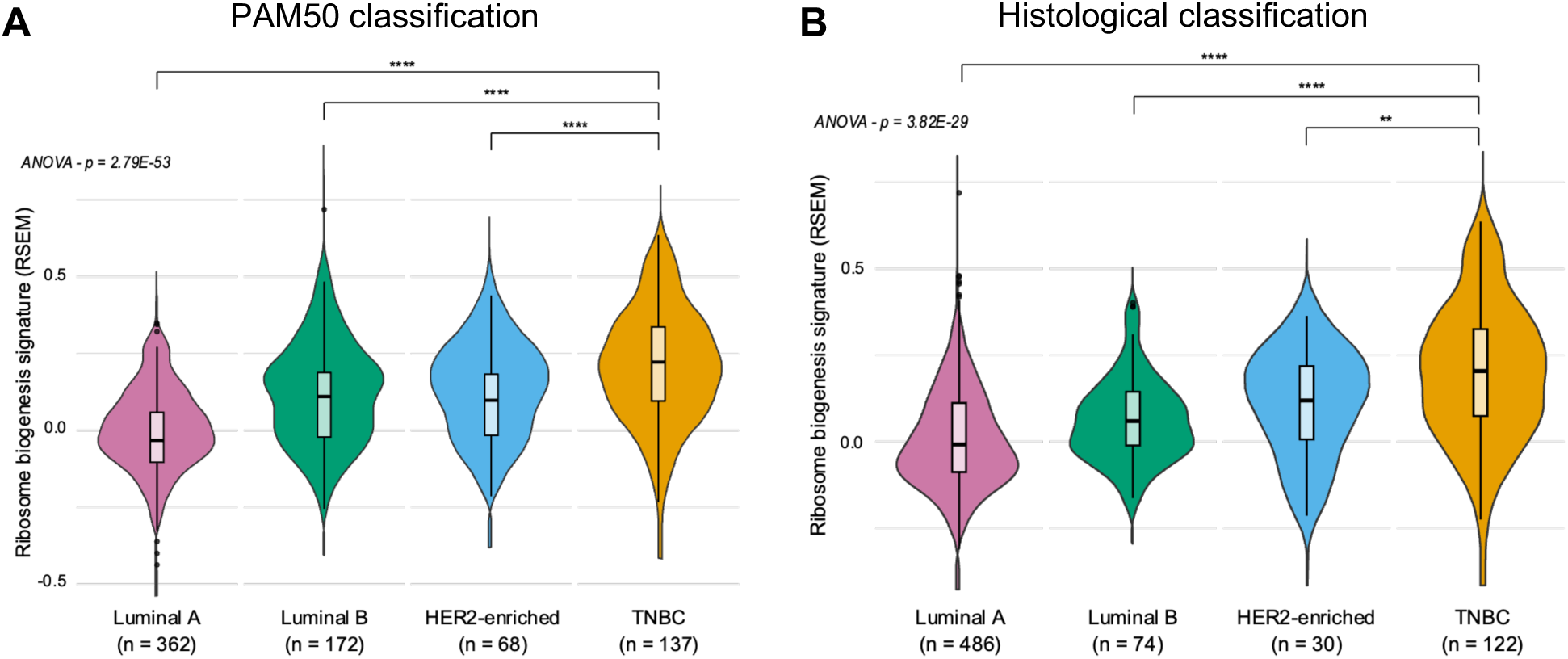
Ribosome biogenesis is higher in TNBC tumors. Comparison of the median expression of ribosome biogenesis genes (240 genes, see Supplementary Table S1) between the different breast cancer subtypes extracted from the UCSC XENA database. The different breast cancer subtypes were defined using either the PAM50 (**A**) or the histological (**B**) classification. n: number of samples; ** p < 0.01, *** p < 0.001 **** p < 0.0001 (ANOVA test, Student test, Mann-Whitney test).

### CX-5461 and BMH-21 display on- and off-target activities in TNBC cells

Given that ribosome biogenesis is overactivated in TNBC, we investigated the impact of its inhibition on TNBC cell lines by blocking RNA Pol I activity, using CX-5461 and BMH-21, two of the most extensively studied RNA Pol I inhibitors. We first determined whether CX-5461 and BMH-21 inhibited RNA Pol I activity in TNBC cells. rRNA synthesis was evaluated by RNA FISH performed using two probes binding to the 5’-ETS and ITS1 regions of the pre-rRNA for the detection of both long- and short-lived pre-rRNA intermediates. In the TNBC cell lines MDA-MB-231 and BT-20, both CX-5461 and BMH-21 efficiently inhibited pre-rRNA synthesis after 4 h of treatment at 1 µM (Figure 2A). Single cell measurement of the RNA FISH signal revealed that BMH-21 was as effective as actinomycin D, used as reference inhibitor of RNA Pol I to inhibit rRNA synthesis, unlike CX-5461 for which inhibition was not complete, as previously described (Figure S1, (7). Furthermore, consistent with its mechanism of action (8), BMH-21 treatment resulted in the destabilization of the RPA-194 RNA Pol I subunit, which decreased by 30% in MDA-MB-231 and 65% in BT-20 cells (Figure 2B). We observed that CX-5461 also reduced RPA-194 by 50% in BT-20, but not in MDA-MB-231, an effect not previously reported (Figure 2B). These data confirmed the high level of activity of CX-5461 and BMH-21 towards RNA Pol I in TNBC cells.

**Figure 2.**
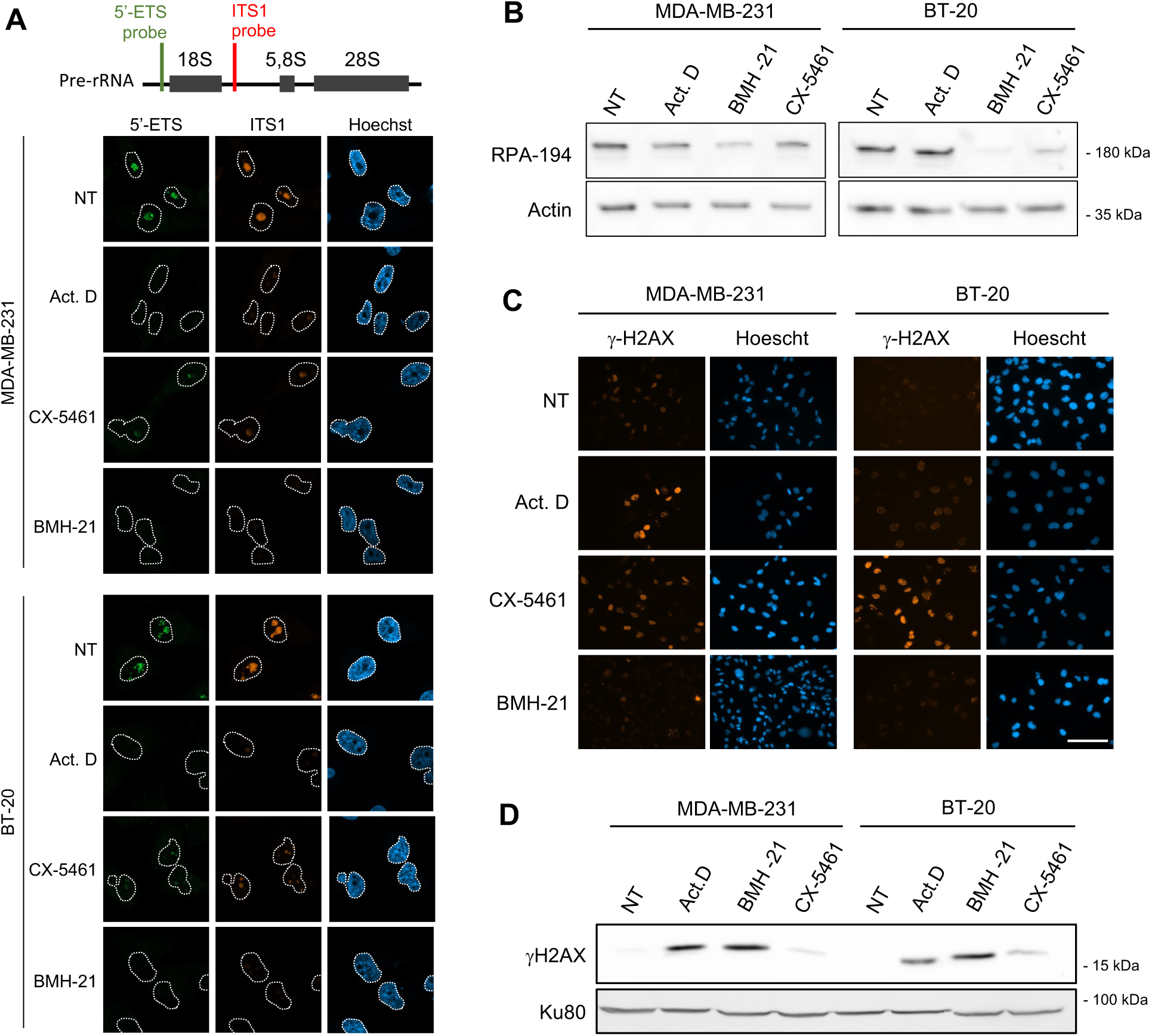
CX-5461 and BMH-21 inhibit RNA Pol I and induce DNA damage in TNBC cells. **A,** Representative images of FISH detection of the pre-rRNA using probes specific to 5’-ETS and ITS1 region in MDA-MB-231 and BT-20 cells, either untreated (NT) or treated with 1 µM CX-5461, 1 µM BMH-21, or 0.05 µg/mL actinomycin D for 4 h. Nuclei were stained with Hoechst. Scale bar: 20 µM. **B,** Immunoblot of RPA-194 protein in MDA-MB-231 and BT-20 cell lines either untreated (NT) or treated with 1 µM CX-5461, 1 µM BMH-21, or 0.05 µg/mL actinomycin D for 4 h. Data representative of 2 independent biological replicates. **C,** Immunofluorescence staining of γ-H2Ax in MDA-MB-231 and BT-20 cells either untreated (NT) or treated with 1 µM CX-5461, 1 µM BMH-21, or 0.05 µg/mL actinomycin D for 24 h. Nuclei were stained with Hoechst. Scale bar: 50 µM. Images are representative of 3 independent biological replicates. **D,** Immunoblot of γ-H2Ax levels in MDA-MB-231 and BT-20 cells lines either untreated (NT) or treated with 1 µM CX-5461, 1 µM BMH-21, or 0.05 µg/mL actinomycin D for 24 h. Data representative of 2 independent biological replicates.

Previous studies have reported conflicting data on possible off-target activity of CX-5461 and BMH-21, especially DNA damage induction, due to their mode of action, binding respectively to G-quadruplex and GC-rich DNA sequences. While DNA damage by BMH-21 remains uncertain, several reports clearly described a genotoxic effect of CX-5461 in some pre-clinical models, likely due to a topoisomerase II inhibitor activity (9,26). Here, DNA damage induction was monitored in TNBC cells by γ-H2AX staining 24 h after treatment with 1 µM of CX-5461 and BMH-21 (Figure 2C and 2D). The level of γ-H2AX staining and the number of its foci, increased after CX-5461 and actinomycin D treatment, but not after exposure to BMH-21 (Figure 2C), a result confirmed by Western blot (Figure 2D). Hence, the RNA Pol I inhibitor CX-5461 induces DNA damage in TNBC cells *in vitro*, while BMH-21 has a very limited impact. Overall, these results demonstrate that RNA Pol I inhibitors CX-5461 and BMH-21 can effectively inhibit ribosome biogenesis in TNBC cells, and that CX-5461 also displays an off-target activity owing to its genotoxic effect, and are consistent with previously published data on cell lines from other cancers (9,26).

### CX-5461 and BMH-21 reduce TNBC cell growth *in vitro*

To evaluate the antitumor activity of the RNA Pol I inhibitors on TNBC cells, the MDA-MB-231 and BT-20 cell lines were treated with CX-5461 and BMH-21 at concentrations ranging from 1-1,000 nM. Both compounds exhibited dose-dependent activities on TNBC cell proliferation (Figures 3A and S2A). IC_50_ values were calculated for each cell line based on cell confluence after 96 h of treatment. The IC_50_ values for CX-5461 in MDA-MB-231 and BT-20 cells were 49.62 nM and 163.5 nM, respectively. For BMH-21, the IC_50_ values were 107.2 nM in MDA-MB-231 and 172.7 nM in BT-20 (Figure 3A). Thus, low concentrations of both compounds induced toxicity, with IC_50_ values in the nanomolar range in both cell lines. We then assessed the effects of CX-5461 and BMH-21 on the clonogenic potential of TNBC cells. Treatments with either compound significantly reduced the number of colonies formed by both cell lines compared to untreated controls at all tested concentrations, notably at 200 nM of CX-5461 and 300 nM BMH-21 (Figure 3B and 3C). Indeed, upon exposure to 200 nM CX-5461, the clone surface area decreased by 60% in BT-20 cells and by 80% in MDA-MB-231 cells, with a 90% reduction observed at 700 nM. BMH-21 also reduced the ability of cells to form clones, with surface areas decreasing by 80-90% at 300 nM, and becoming negligible at higher concentrations (1,000 nM) (Figure 3C). Overall, the two compounds drastically impaired TNBC cell viability when cultured as a monolayer.

**Figure 3.**
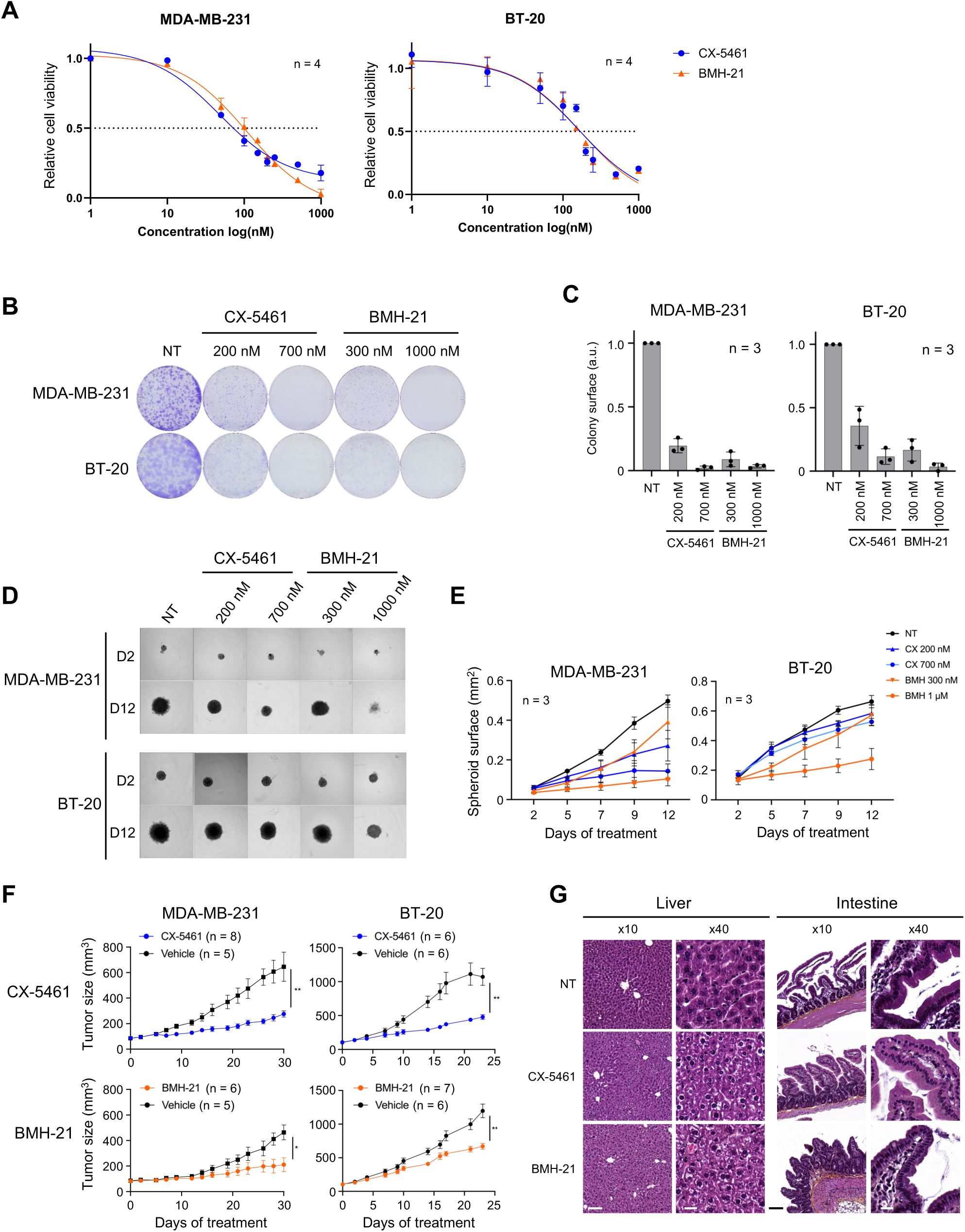
Inhibition of RNA Pol I reduces tumorigenesis *in vitro* and *in vitro* in TNBC. **A,** Relative cell proliferation of MDA-MB-231 and BT-20 cells following treatment with 1-1,000 nM CX-5461 and BMH-21. Error bars correspond to the s.d. of 4 biological replicates. **B-C,** Colony formation assay on MDA-MB-231 and BT-20 cells, either untreated (NT) or treated with CX-5461 or BMH-21, at the indicated concentration for 10 days. The surface of the colony was quantified (**C**) from the images in (**B**). The result is representative of 3 independent biological replicates. The data are presented as mean ± s.d. **D,** Spheroid growth of MDA-MB-231 and BT-20 cells either untreated (NT) or treated with CX-5461 or BMH-21 at the indicated concentrations. Images shown were collected 2- or 12-days post-treatment. **E,** the size of the spheroids was quantified from the images in (**D**). The results are mean ± SEM from 3 technical replicates, and are representative of 3 biological replicates. **F,** Growth of tumor xenografts of MDA-MB-231 and BT-20 cells over 30 days, in mice treated with CX-5461 or BMH-21 at 50 mg/kg, or vehicle at day 0. Quantification of tumor volume is presented as mean ± SEM (n = 5). * p < 0.05, ** p < 0.01 (Student t-test) **G,** Hematoxyline-eosine staining of liver and intestine tissue sections, from mice treated 23 days with CX-5461 or BMH-21 or from control animals (Vehicle). Images were collected using a x10 and x40 objective. Scale: at x10 = 100 µm; at x40 = 20 µm. Representative images of 3 animals.

We next evaluated the effects of CX-5461 and BMH-21 on the ability of TNBC cells to grow in 3D as spheroids. TNBC spheroids were treated as indicated 2 days post-formation and were monitored every 2 to 3 days by microscopic observation, and the surface area of each spheroid was measured (Figure 3D and 3E). Treated spheroids were smaller compared to untreated controls, with a stronger effect observed at higher concentrations. For both cell lines, spheroid growth displayed a reduction as early as day 5 post-treatment in a dose-dependent manner. Exposure to 200 nM or 700 nM CX-5461 for 12 days resulted in a 1.8-fold and 3.4-fold reduction in the size of MDA-MB-231 spheroids, respectively, while only a slight decrease was observed for BT-20 spheroids (Figure 3D and 3E). Conversely, BMH-21 similarly impacted the growth of spheroids of both cell lines, with a limited impact at 300 nM, and a 5-fold reduction on MDA-MB-231 spheroids and 2.4-fold reduction on BT-20 spheroids at 1,000 nM, by day 12 (Figure 3D and 3E). Interestingly, the sensitivity of cells grown in 3D was similar to that of cells grown in monolayer. Overall, our results show that RNA Pol I inhibitors CX-5461 and BMH-21 reduce TNBC cell growth *in vitro*, even at nanomolar concentrations.

### CX-5461 and BMH-21 reduce tumor growth in human TNBC xenografts

The antitumor activity of CX-5461 and BMH-21 was then investigated in MDA-MB-231 and BT-20 xenograft models, upon subcutaneous injection into athymic nude mice. CX-5461 was administered orally at 50 mg/kg three times a week, while BMH-21 was given intraperitoneally at 50 mg/kg five times a week, as previously described (7,8). In these models, CX-5461 treatment significantly reduced tumor growth by 2 to 3-fold compared to the control group (from 645 mm^3^ to 276 mm^3^ in MDA-MB-231, and from 1070 mm^3^ to 477 mm^3^ in BT-20) (Figure 3F), and BMH-21 also significantly reduced tumor size, though to a lesser extent (from 463 mm^3^ to 210 mm^3^ in MDA-MB-231, and from 1093 mm^3^ to 669 mm^3^ in BT-20) (Figure 3F). Given the genotoxic activity of CX-5461 *in vitro* (Figure 2C-D), we wondered whether it may have a similar effect in xenografts. Consistent with our observation in cultured cells, CX-5461 treatment, but not BMH-21, induced H2AX phosphorylation in the xenografts (Figure S2B, demonstrating that, as in other cancers (9,26), both RNA Pol I inhibitors impair TNBC tumor growth *in vivo*, and that CX-5461 alone exhibits an off-target activity.

Importantly, given that ribosome biogenesis occurs in all cells, it was crucial for us to demonstrate the selective effects of RNA Pol I inhibitors on tumors. As previously described, we observed no impact on the body weight of our xenograft models treated with both compounds (Figure S2C (7,8)). To further evaluate their potential toxicity, we performed a histological analysis of several organs, including the brain, heart, lungs, liver, kidneys, pancreas, stomach, intestine, and colon. No major lesions were detected in the tissues of mice treated with BMH-21 or CX-5461. However, some moderate histological alterations were noted. In the intestine, villous atrophy was observed in BMH-21-treated mice, while no changes were detected in the intestines of control or CX-5461-treated mice. Additionally, a diffuse oedema was observed in the colon of both BMH-21- and CX-5461-treated mice compared to controls. In the liver, hepatocytes with clear cytoplasm were diffusely present in BMH-21-treated mice, whereas this effect was localized to the periportal region in CX-5461-treated mice (Figure 3G). Overall, these findings suggest minimal acute toxicity of CX-5461 and BMH-21 following short-term treatment in mice. However, they highlight the need for caution regarding potential adverse effects associated with long-term use.

### CX-5461 and BMH-21 induce cell cycle arrest but no cell death *in vitro* and *in vivo*

Ribosome biogenesis inhibition is known to trigger ribosomal stress, which can lead to cell death and/or cell cycle arrest. We thus explored these two cell responses in MDA-MB-231 and BT-20 cell lines, through DRAQ7 and cleaved caspase-3 staining. Following three days of treatment, no evidence of cell death was observed in cells treated with CX-5461 or BMH-21, while accumulating dead cells was observed upon actinomycin D treatment in both MDA-MB-231 and BT-20 cell lines (Figure S3A). Consistently, in the xenograft tumors, immunohistochemical staining of cleaved caspase 3 indicated that apoptosis was not triggered in response to CX-5461 and BMH-21 treatment (Figure S3B). Thus, CX-5461 and BMH-21 do not induce cell death in these TNBC cell lines both *in vitro* and *in vivo*.

Next, we evaluated the effects of CX-5461 and BMH-21 on cell cycle progression. Treatment with CX-5461 (200 and 700 nM) and BMH-21 (300 and 1,000 nM) for 18 h resulted in distinct alterations in cell cycle progression (Figures 4A and S3C-S3D). For CX-5461, treatment led to a significant accumulation in the G2/M phase in both MDA-MB-231 and BT-20 cells, with higher concentrations also causing an accumulation in the S phase and a reduction in the G1 phase in MDA-MB-231. CX-5461 treatment thus promotes S-phase defect and G2/M checkpoint arrest. For BMH-21, the effects were cell-dependent, with the induction of a significant accumulation of MDA-MB-231 cells in the G1 phase, along with a decrease in the S phase, indicative of a G1/S checkpoint arrest. In contrast, BT-20 displayed no clear alteration of cell cycle despite the growth defect observed (Figures 3A, 4A and S3C). We further explored the different effects on the cell cycle status induced by the two RNA Pol I inhibitors, by focusing on the M phase through phospho-histone H3 (Ser10) staining (Figure S3D). Compared to CX-5461, BMH-21 treatment induced a marked decrease in phospho-H3, in particular in BT-20 cells, indicating that BMH-21 prevented mitosis (Figure S3C). To determine the relevance of these observations in *in vivo* models, we performed a concomitant IHC analysis of proliferation (Ki67 marker) and mitosis status (phospho-H3 staining) in xenografted tumors. We observed no difference in the Ki67 signal in the proliferative area of the tumor, for both cell lines and both treatments (Figure 4B). In contrast, a decrease in phospho-H3 (Ser10) signal was observed in MDA-MB-231 and BT-20 xenografts upon treatment with CX-5461 (Figure 4B), supporting the cell-type dependent effects of these RNA Pol I inhibitors on cell cycle arrest *in vitro* and *in vivo*.

**Figure 4.**
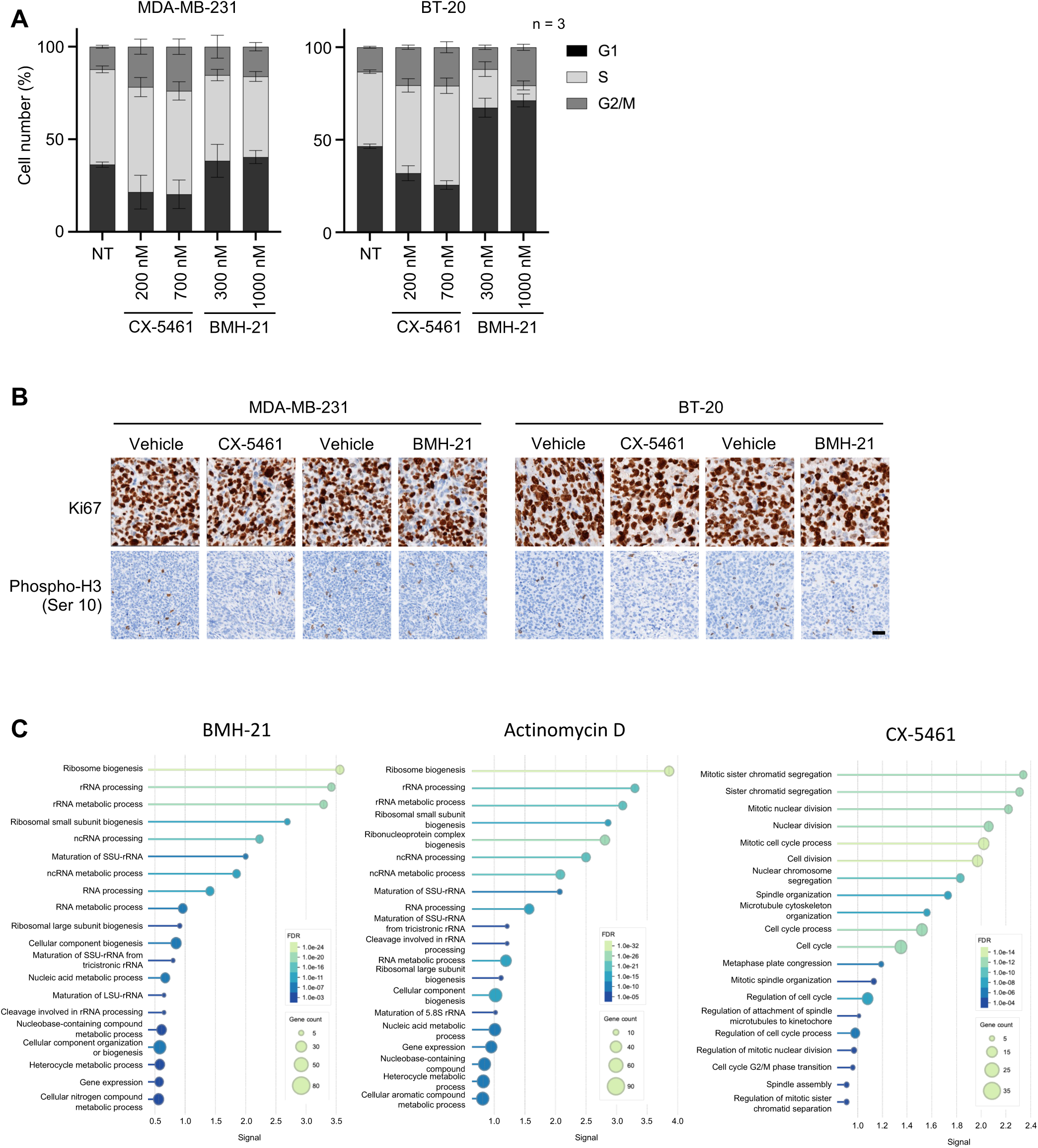
Impact of CX-5461 and BMH-21 on cell cycle and ribosome biogenesis in TNBC cells *in vitro.* **A,** Proportion of cells in G1, S, and G2 phases of the cell cycle of MDA-MB-231 and BT-20 cells, either untreated (NT) or treated with CX-5461 or BMH-21, at the indicated concentrations or with actinomycin D at 0.05 µg/mL for 18h. Error bars correspond to the s.d. from three technical replicates. Results are representative of six biological replicates. **B,** IHC staining of the proliferation marker Ki67 and mitosis marker phospho-Histone H3 (Ser 10), on sections of xenograft tumors, from the experiment shown in Figure 3F. Images are representative of 5 biological replicates. Scale bar: 50 µm. **C,** Gene ontology analysis performed on proteomic data obtained on whole cell extracts of MDA-MB-231 cells treated with CX-5461, BMH-21 or actinomycin D for 18h (see Table S6). The graph shows the 20 most represented GO terms (see Table S7) among the differentially expressed genes compared to untreated cells. The data were obtained from 5 biological replicates.

To better understand the TNBC cell response to these inhibitors, we then performed a label-free quantitative proteomics analysis on whole cell extracts of MDA-MB-231 cells treated with CX-5461, BMH-21 or actinomycin D as a reference treatment. The analysis identified and quantified 4,513 proteins. To evaluate changes in protein levels, pairwise comparisons based on mass spectrometry (MS) peak intensity values were performed for each protein between treated and untreated condition. Each treatment induced a notable alteration in the cell proteome (Figure S4 and Table S6), with significant alterations of the level of 277 proteins in actinomycin D treated cells, 127 proteins in CX-5461 treated cells and 156 proteins in BMH-21 treated cells (Figure S4). Interestingly, among the differentially expressed proteins following BMH-21 and actinomycin D treatment, a majority were down-regulated (76.9 % and 81.2 respectively), whereas following CX-5461 treatment a minority of proteins were down-regulated (34.0 %), reflecting a different impact on the cell gene expression (Figure S4). We then performed a Gene Ontology (GO) analysis in order to identify the most altered cellular functions (Figure 4C and supplementary table S7). The most represented GO Terms were very different among the three conditions, actinomycin D and BMH-21 treatments being associated with ribosome biogenesis and nucleic acid-related cellular functions, while CX-5461 treatment was associated with cell division-related cellular functions (Figure 4C). These findings further characterize BMH-21 as a *bona fide* ribosome biogenesis inhibitor, and support that CX-5461 does not act as a ribosome biogenesis inhibitor but rather as a cell division inhibitor. Collectively, these findings emphasize the potential of ribosome biogenesis as a therapeutic target in TNBC. However, they also highlight the need for developing more specific strategies to inhibit ribosome biogenesis, while minimizing off-target effect.

### FBL expression is higher in TNBC compared to other breast cancer subtypes

Having shown that ribosome biogenesis is a druggable vulnerability of TNBC, we next sought to identify other ribosome biogenesis factors that could be inhibited for therapeutic purposes. Once the rRNA precursor is synthesized by RNA Pol I, it undergoes several essential cleavages that release the 18S, 5.8S and 28S (27). C/D box snoRNP containing the U3 snoRNA is required for a cleavage step within the 5’-ETS that is essential for rRNA maturation. It has been reported that the knock-down of U3 snoRNA decreases the growth of lung and luminal breast cancer cells, and prevents tumor initiation of lung cancer cell xenografts (15). In addition, we have previously shown that the knock-down of FBL, a central component of the C/D snoRNP, inhibits the U3 snoRNA-driven cleavage step within the 5’-ETS region of the pre-rRNA (16). We thus explored FBL as a potential therapeutic vulnerability in TNBC. We previously demonstrated that elevated FBL mRNA levels are associated with poor survival in breast cancer patients, although this analysis did not focus on specific breast cancer subtypes (20). To determine whether the C/D snoRNP complex signature differs in TNBC, we compared this signature (*i.e.*, FBL, NOP56, NOP58, NHP2L1) in different breast cancer subtypes using the BRCA-TCGA dataset as described above for the ribosome biogenesis signature. The C/D snoRNP complex signature was significantly higher in TNBC than in other breast cancer subtypes, using either the PAM50 classification (ANOVA, p = 9.06 x 10^−51^, Figure 5A) or the histological classification (Figure S5A). Importantly, the expression level of *FBL* itself was higher in the TNBC subtype compared to other breast cancer subtypes (ANOVA, p = 8.69 x 10^−41^, Figure 5B and Figure S5B).

**Figure 5.**
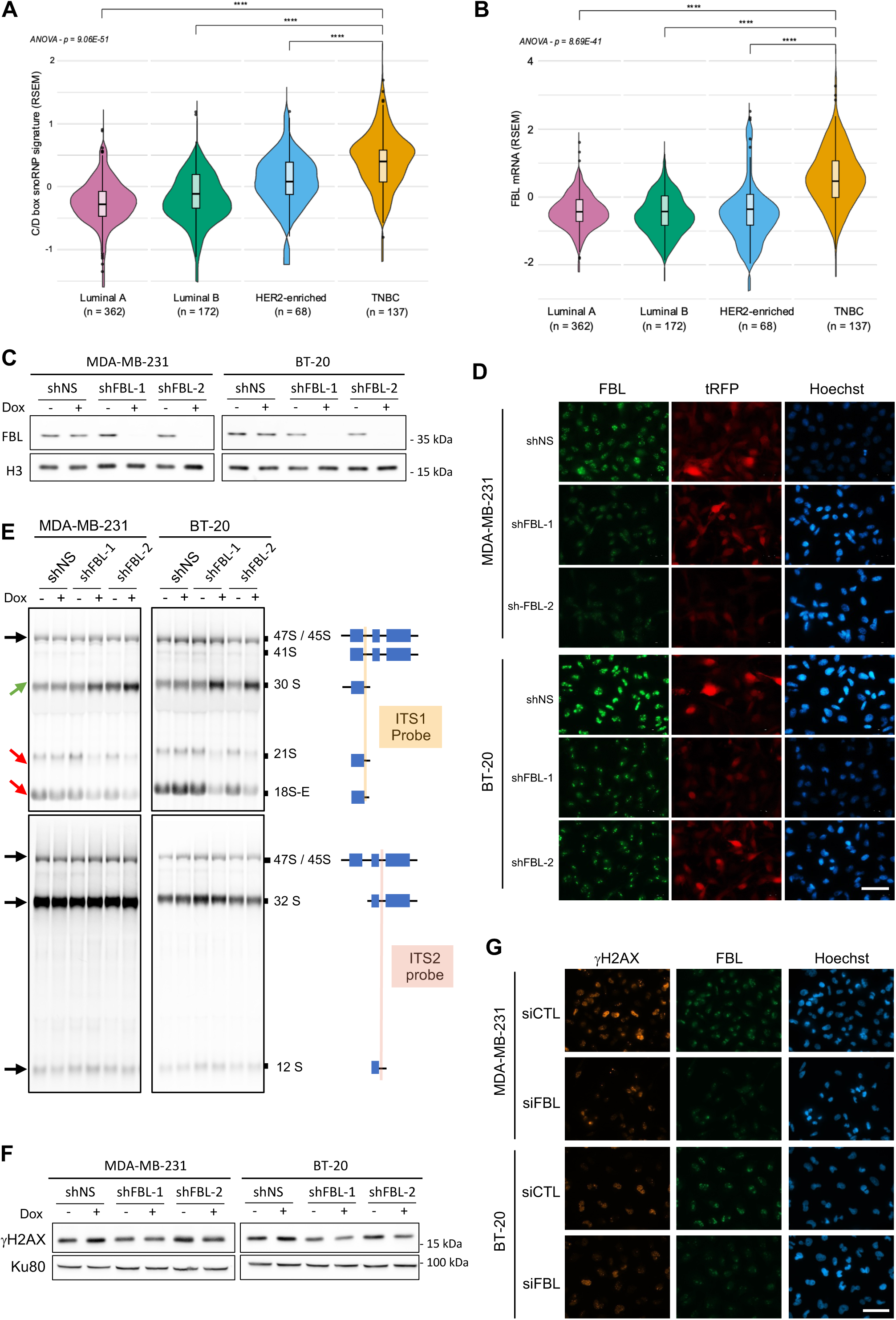
FBL knock-down inhibits maturation of pre-rRNA whithout inducing DNA damage. **A-B,** Comparison of the median expression of mRNAs of C/D box snoRNP protein genes (**A**) (FBL, NOP56, NOP58, NHP2L1) or of FBL (**B**), between each breast cancer subtypes. Normalized transcriptomic data were extracted from the UCSC XENA database. The breast cancer subtypes were defined using the PAM50 classification. n: number of patients, **** p < 0.0001 (ANOVA test, Mann-Whitney test). **C,** FBL protein levels analysed by Western blot in MDA-MB-231 and BT-20 cell lines after 96-h doxycycline-mediated shRNA induction. The cell lines express either a control non-silencing shRNA (shNS) or a shRNA-targeting FBL (FBL-1 and FBL-2). Data are representative of 3 independent replicates. **D,** Immunofluorescence staining of FBL in MDA-MB-231 and BT-20 cell lines after 96-h doxycycline-mediated shRNA induction (shFBL-1, FBL-2, and shNS). The tRFP protein is co-expressed with the shRNA and is used as a marker of doxycycline induction. The images are representative of 3 biological replicates. Scale bar: 50 µm. **E,** Northern blot analysis of pre-rRNA processing in MDA-MB-231 and BT-20 cells after 96-h doxycycline-mediated shRNA induction (shFBL-1, FBL-2, and shNS). The positions of the probes used and the detected pre-rRNA species are indicated on the right. Arrows on the left show the trends of each pre-rRNA species in shRNA-induced cells relative to non-induced cells. Data representative of two biological replicates. **F,** Western blot analysis of γ-H2Ax protein levels in MDA-MB-231 and BT-20 cell lines after 96-h doxycycline-mediated shRNA induction (shFBL-1, FBL-2, and shNS). Data representative of two biological replicates. **G,** Immunofluorescence staining of γ-H2Ax and FBL in MDA-MB-231 and BT-20 cells transfected with a FBL siRNA or a control siRNA. Scale bar: 50 µm.

### FBL knock-down inhibits pre-rRNA maturation and cell growth *in vitro* without inducing DNA damage in TNBC cells

To assess the impact of FBL knock-down on TNBC cell lines, we established MDA-MB-231 and BT20-based stable cell lines, expressing either two independent FBL shRNAs (*i.e.*, shFBL-1 and shFBL-2) or a negative control shRNA (shNS), in an inducible manner. Effective knock-down of FBL expression was confirmed at the mRNA level by RT-qPCR (Figure S5C) and at the protein level by Western blot (Figure 5C), showing more than 90% reduction in both mRNA and protein levels. Immunofluorescence imaging revealed a homogeneous decrease in FBL signal within the cell population (Figure 5D). Of note, we observed a significant reduction in FBL levels from 2 days post-induction onwards in both cell lines (Figure S5D), whereas doxycycline and non-specific shRNA (*i.e.*, shNS) had no impact on its expression. To determine whether FBL knock-down impairs pre-rRNA processing in TNBC cells, Northern blot analysis was performed to examine different cleavage stages of the 18S and 28S/5.8S rRNA processing pathways (Figure 5E, ITS1 probe and ITS2 probe, respectively). Following 96 h of FBL knockdown, 18S rRNA processing was significantly inhibited, as illustrated by the accumulation of early-stage intermediate species (*i.e.,* 30S pre-rRNA) and a decrease in late intermediate species (*i.e.,* 21S and 18S-E pre-rRNAs). In contrast, no impact on the 28S / 5.8S processing pathway was observed (Figure 5E, ITS2), as previously reported in HeLa cells (16).

Because of the off-target activity observed with the RNA Pol I inhibitors, we wondered whether FBL knock-down induces DNA damage. Our analysis revealed that FBL reduction did not cause detectable DNA damage as assessed by Western blot analysis of γ-H2AX (Figure 5F) and by immunostaining (Figure 5G and S5E). These findings were consistent across both shRNA-based (Figure 5F) and siRNA-based knock-down approaches (Figure 5G and S5E). Hence, FBL knock-down impairs ribosome biogenesis by blocking the 5’-ETS pre-rRNA maturation, without inducing DNA damage, in our TNBC cellular models.

Given the inhibitory effect of FBL knock-down on TNBC cell ribosome biogenesis, we determined whether its targeting could be anti-tumoral using various *in vitro* assays. Monitoring cell growth in real-time revealed that FBL knock-down significantly reduced cell proliferation compared to controls (shNS and non-treated cells) (Figure 6A). Even after 250 h (10 days), FBL knock-down cells reached no more than 40% confluence, whereas non-treated cells achieved 100% confluence within 100 h (4 days) (Figure 6A). A similar impact was observed on colony formation (Figure 6B) and on the surface area of the colonies, which decreased by 80% to 95% in FBL knock-down cells compared to control conditions (Figure 6C). We then determined whether FBL knock-down also impaired the proliferation of cells grown in 3D as spheroids, by inducing shRNA expression with doxycycline for 12 days. Upon expression of either one of the FBL shRNAs, the spheroids were significantly smaller than non-treated control and shNS spheroids (Figure 6D and 6E). After 12 days, shFBL-1 reduced the size of MDA-MB-231 spheroid by 1.8-fold, while shFBL-2 caused a 1.6-fold reduction. In BT-20 cells, shFBL-1 reduced spheroid size by 4-fold, and shFBL-2 by 3.3-fold (Figure 6E). Additionally, we noticed that shFBL-expressing spheroids exhibited an altered structure, characterized by a lower cell density and irregular edges, a phenotype that was more pronounced on MDA-MB-231 spheroids than on BT-20 spheroids (Figure 6D). In conclusion, our data show that decreasing FBL expression inhibits cell growth of TNBC cells *in vitro*.

**Figure 6.**
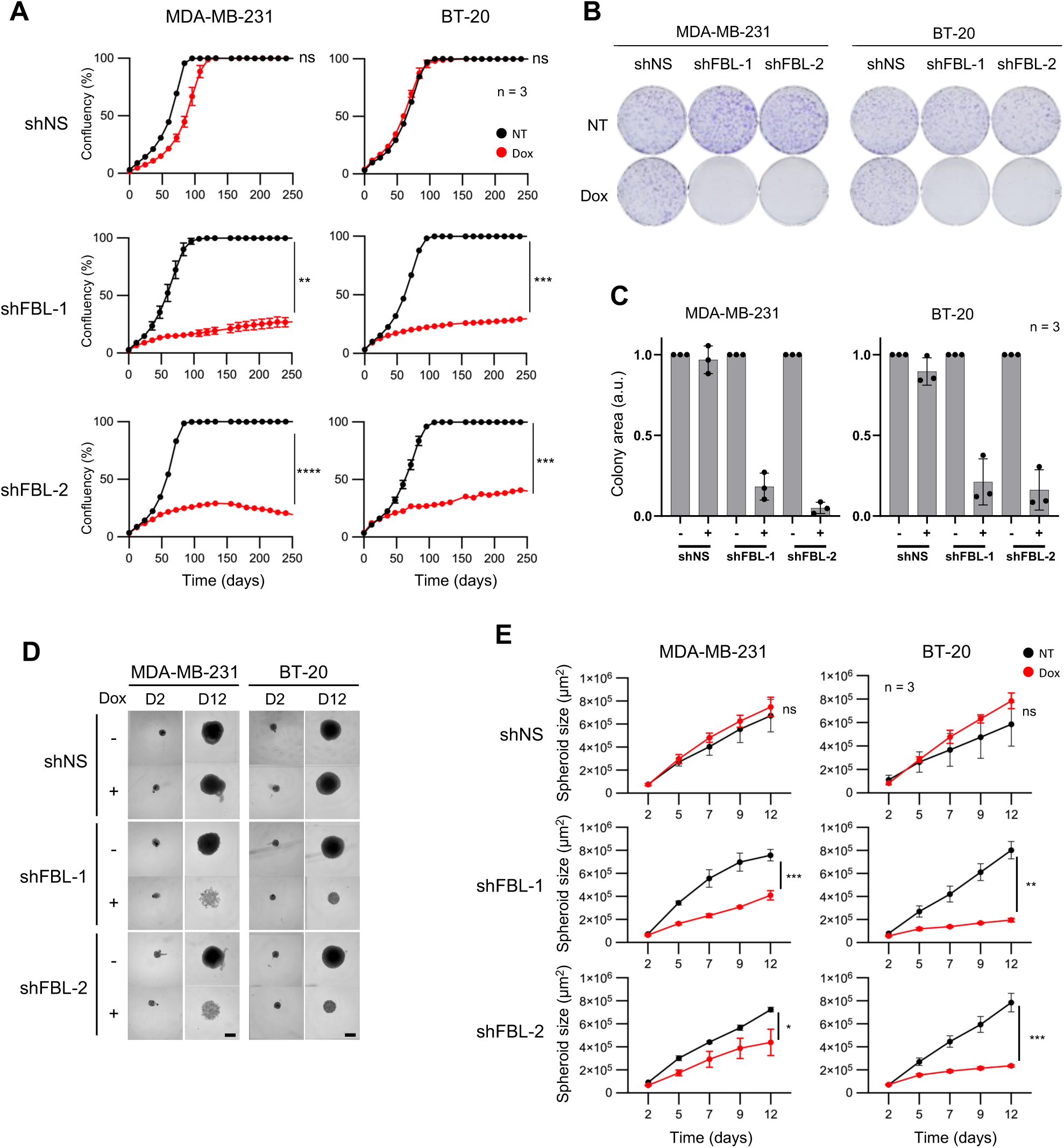
Knock-down of FBL exerts an antitumoral activity on TNBC cells. **A,** Real time monitoring of the proliferation of MDA-MB-231 and BT-20 cell lines following induction of shRNA expression with doxycycline (Dox), compared to non-induced cells (NT). Data represent mean ± s.d. of three technical replicates. Student t-test. Data are representative of three biological replicates. **B-C,** Colony formation assay with MDA-MB-231 and BT-20 cell lines upon doxycycline induction of shRNA (shFBL-1, FBL-2, and shNS). Images in (**B**) are representative of three independent experiments at day 12. Quantification of colony surface area (**C**) from three biological replicates experiments. The surface area was normalised against the untreated condition for each doxycycline-treated condition. Data represent mean values ± s.d. and individual measurements (dots) **D-E,** Spheroid formation with MDA-MB-231 and BT-20 cells upon doxycycline induction (shFBL-1, FBL-2, and shNS). Phase contrast images were collected every 2 or 3 days, and representative images of spheroids at day 2 and day 12 are shown (**D**). Spheroid surface was quantified over time (**E**), with (Dox) or without (NT) doxycycline induction. Data represent the mean ± s.d. of three biological replicates.

### FBL knock-down induces cell cycle defects but no cell death in TNBC cells *in vitro*

We next characterized the mechanisms underlying the anti-tumoral effects of FBL knock-down. After 96 h of FBL knock-down, no evidence of cell death was observed (below 0.5%), while actinomycin D treatment led to 20 to 40% cell death, as reported above (Figure S3A and Figure 7A). Analysis of apoptotic markers by Western blot, including caspase 3 and PARP-1, revealed no significant difference in the expression or cleavage of these proteins following FBL knock-down (Figure 7B). These findings indicated that apoptosis was not triggered, and were consistent across both shRNA and both cell lines, and were further confirmed through the transfection of FBL siRNA in both TNBC cell lines (Figure S6A). Thus, as observed with RNA Pol I inhibitors, ribosome inhibition by FBL knock-down did not trigger cell death in our TNBC cellular models.

**Figure 7.**
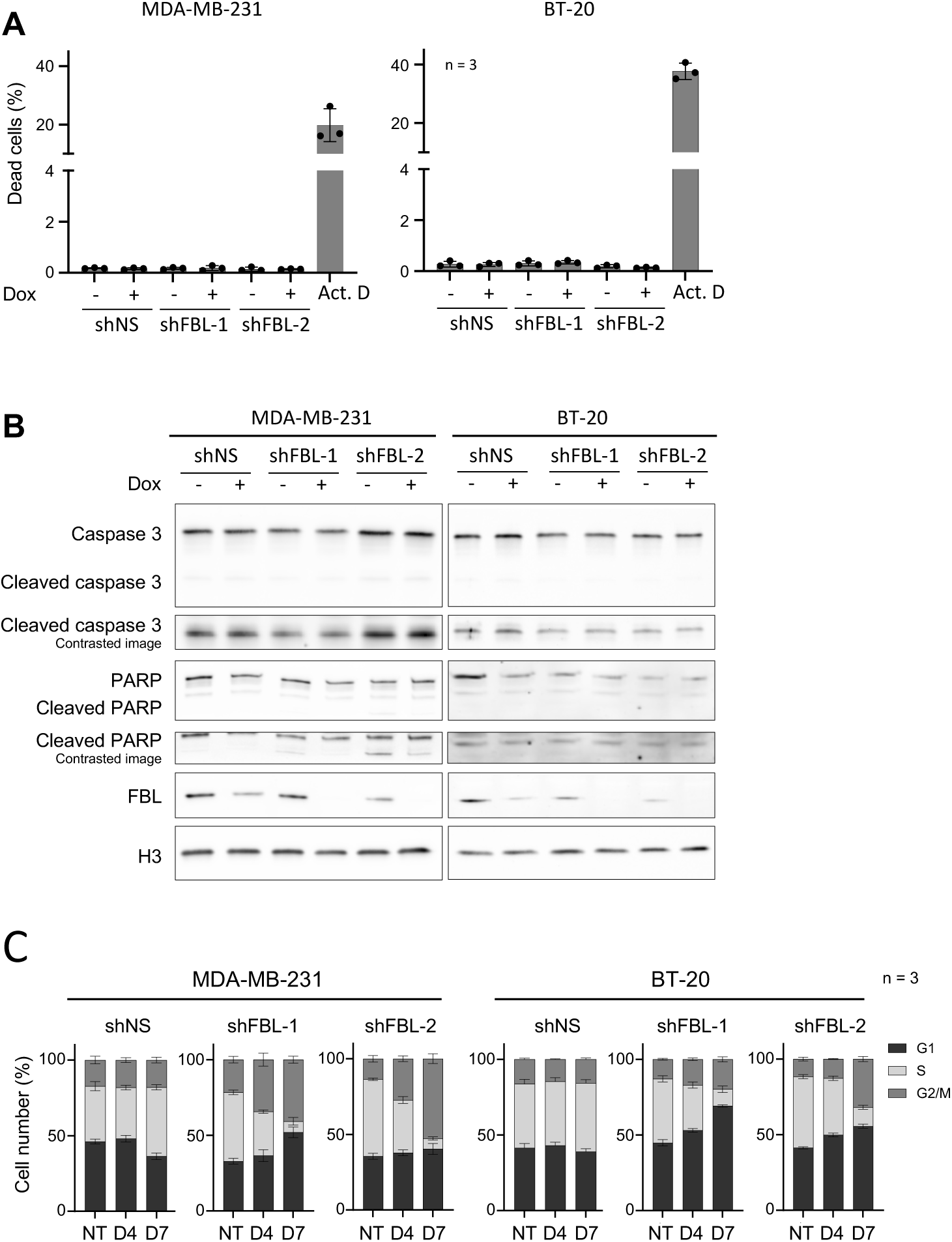
Knock-down of FBL induces cell cycle defect but no cell death. **A,** Cell death analysis on MDA-MB-231 and BT-20 cells following 4 days of doxycycline induction (shFBL-1, FBL-2, and shNS) or 18 h treatment with actinomycin D, measured by DRAQ7 staining and fluorescence imaging. Data represent mean values ± s.d. of the percentage of dead cells from a technical triplicate, and are representative of two independent biological replicates. **B,** Western blot showing full length and cleaved caspase 3 and PARP proteins analysed in MDA-MB-231 and BT-20 cell lines following doxycycline-mediated shRNA induction for 72 h. Contrasted images are shown for the detection of cleaved forms. FBL levels are shown to confirm the efficacy of its knock-down. Data representative of three biological replicates. **C,** Proportion of cells in G1, S, and G2 phases of the cell cycle of MDA-MB-231 and BT-20 cells, following doxycycline-mediated expression of shRNA (shFBL-1, FBL-2, and shNS) for 4 or 7 days. Data are mean values +/− s.d. of three technical replicates. Results are representative data of three biological replicates.

In contrast, we observed that FBL knock-down impaired cell cycle progression, after either four days (D4) or seven days (D7) of shRNA induction (Figure 7C and S6B). In both cell lines and for both FBL shRNAs, we observed an increase in G2/M phase and a strong decrease in S-phase. A more pronounced effect was observed in cells treated for seven days (Figure 7C and Figure S6B). Our results highlight that inhibiting rRNA maturation leads to cell cycle arrest in TNBC, an effect that appears to be more pronounced than the one triggered by CX-5461 and BMH-21 RNA Pol I inhibitors.

### FBL knock-down induces tumor growth defects in TNBC xenografts

Finally, we assessed whether the *in vitro* anti-tumoral effect resulting from FBL knock-down could be replicated in an *in vivo* xenograft model. The MDA-MB-231 cells expressing shFBL-1, shFBL-2 and shNS were subcutaneously xenografted into athymic nude mice. Once tumors were established, shRNA expression was induced by providing the mice with 1 mg/mL doxycycline in their drinking water (Figure 8A and Figure S7). In the MDA-MB-231 model, tumors with FBL knock-down exhibited a significantly reduced growth compared to both the vehicle control and shNS groups, starting 6 to 8 days post-induction (Figure 8A). Tumor growth was negligible in the shFBL-1 model and strongly reduced in the shFBL-2 model.

**Figure 8.**
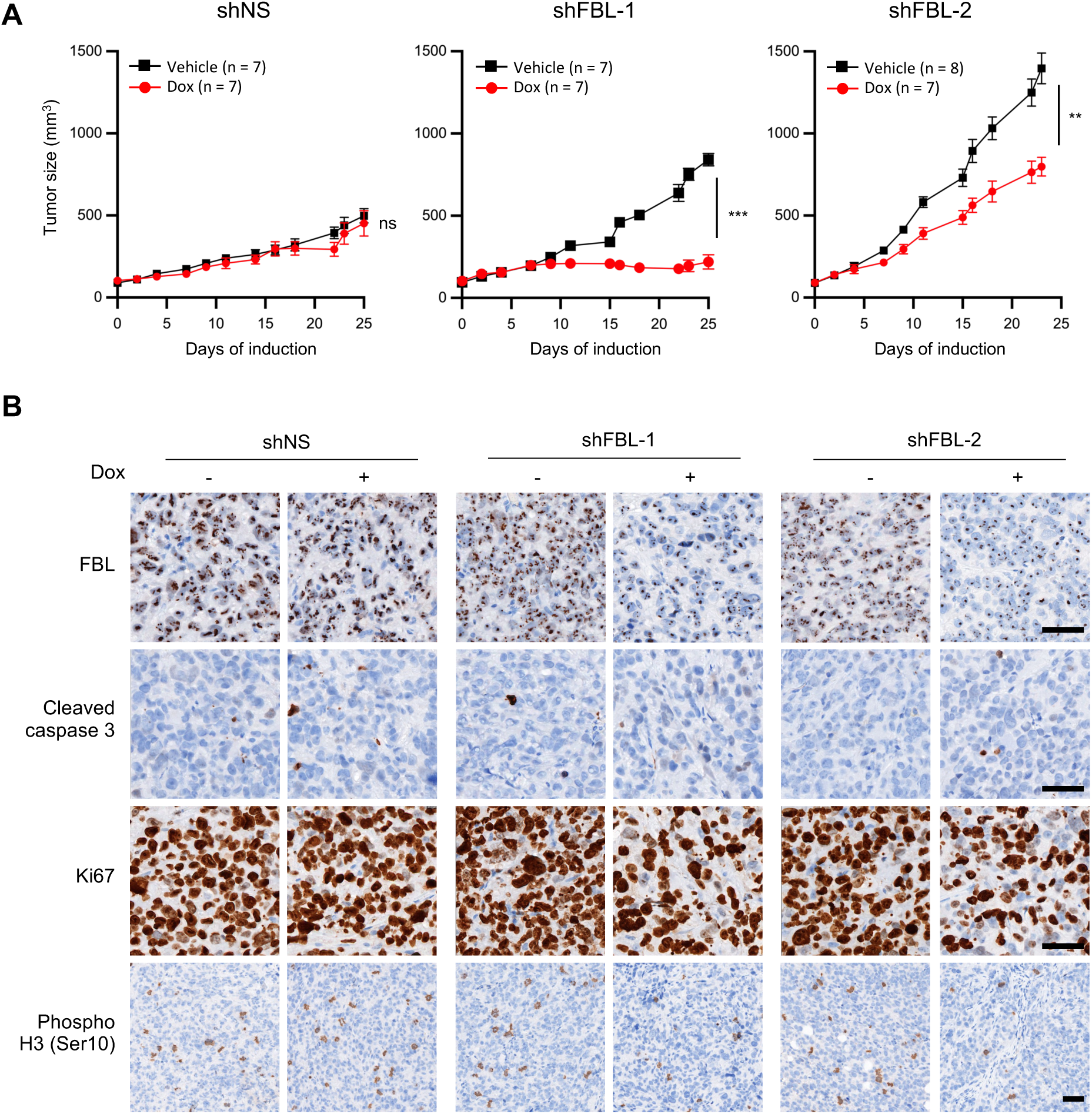
FBL knock-down suppresses tumorigenicity *in vivo* in TNBC. **A,** Growth of tumor xenografts of MDA-MB-231 cell lines expressing either shFBL-1, shFBL-2, or shNS, upon doxycycline induction. Doxycycline was administered via drinking water for 25 days. Data are mean ± SEM. ** p < 0.01, *** p < 0.001 (Student t-test). **B,** Detection of FBL, cleaved caspase 3, Ki67 and phospho-histone H3 (Ser10) by IHC on tissue sections of MDA-MB-231 xenografted tumors from the experiment in (**A**). Images are representative of 5 xenografted tumors. Scale bar: 50 µm.

Immunohistochemistry confirmed a marked reduction in FBL expression only in the dox-treated shFBL xenograft, and neither in the shNS xenograft nor in the non-treated controls (Figure 8B). Importantly, there was no significant weight loss observed in any of the mice throughout the experiment (Figure S7). Thus, FBL knock-down reduces growth of xenografted MDA-MB-231 cells. To further characterize the impact of FBL knock-down on xenografted cells, we analysed apoptosis and proliferation markers by IHC (Figure 8B). Consistent with the *in vitro* data, no cleaved caspase 3 was detected, supporting the conclusion that FBL knock-down does not trigger apoptosis in these cells. Additionally, Ki67 and phospho-Histone H3 (Ser10) signals were strongly reduced upon FBL shRNA expression (Figure 8B), corroborating the *in vitro* findings that FBL knock-down impairs cell cycle progression.

Together, these results demonstrate that inhibition of rRNA maturation through FBL depletion exerts anti-tumoral effects on TNBC xenografts *in vivo* by promoting cell cycle arrest.

## Discussion

Given the poor response of TNBC patients to current standard-of-care treatments and their relapse, novel therapeutic strategies are urgently needed. The anticancer effects of ribosome biogenesis inhibition are well-supported by preclinical and clinical studies across various solid and haematological cancers. In this study, we demonstrate that ribosome biogenesis is an actionable vulnerability in TNBC, using two different strategies to inhibit ribosome production: suppression of rRNA synthesis and disruption of rRNA maturation.

Alterations in ribosome biogenesis have been reported in multiple cancer types, including TNBC, where the ribosome biogenesis machinery and RNA Pol I regulatory pathways are frequently dysregulated (20,24,28). Using transcriptomic data from The Cancer Genome Atlas (TCGA) and a ribosome biogenesis signature comprising 240 genes, we now provide evidence that ribosome biogenesis activity is more elevated in TNBC than in other breast cancer subtypes. Notably, this heightened ribosome biogenesis activity is summarized by a single key factor: the rRNA maturation factor FBL. Our findings reveal that FBL expression is markedly higher in TNBC than in other breast cancer subtypes, providing direct evidence of its link to this aggressive tumor type. TNBC thus represents an additional aggressive cancer subtype characterized by pronounced overactivation of ribosome biogenesis. The abnormal overactivation of ribosome biogenesis in TNBC provides one possible explanation for the sensitivity of TNBC cells to ribosome biogenesis inhibition that we observed in this study. Further studies will be required to extend our observations across additional cell line models that represent the full heterogeneity of the TNBC subtype.

Since the early 2010s, there has been a renewed interest in ribosome biogenesis as a therapeutic target with the discovery and exploitation of the new generation of RNA Pol I inhibitors CX-3543, CX-5461 and BMH-21(Bywater et al., 2012; Drygin et al., 2009; Peltonen et al., 2014). These compounds showed anti-tumoral activity on a wide range of pre-clinical models of haematological and solid tumors, including lymphoma, melanoma, osteosarcoma, and prostate and ovarian cancers (7,8,29–32). Our data demonstrate that the RNA Pol I inhibitors CX-5461 and BMH-21 exhibit potent anti-tumoral activity in pre-clinical models of TNBC both *in vitro* and *in vivo*. In TNBC cells, these inhibitors effectively block pre-rRNA production, thereby disrupting ribosome biogenesis. However, the mechanism of action of both compounds relies on a direct interaction with DNA, particularly G4 quadruplexes and GC-rich sequences, which increases the risk of DNA-based off-target effects. While such off-target activity has herein been established for CX-5461 in TNBC and previously for other cancer types (9,26), a detailed investigation into DNA-related off-target effects of BMH-21 remains to be conducted. In this study, we show that CX-5461, as actinomycin D, induced significant DNA damage and the accumulation of cells in S-phase, which indicate that cells undergo replication-stress, an observation supported by a previous study showing that CX-5461 acts as a Topoisomerase poison (26). Consistently, CX-5461 displayed some toxicity in humans with frequent side effects (nausea, headaches), and significant phototoxicity (32,33). Collectively, these data highlight the need for safer and more specific strategies to target ribosome biogenesis. To this end, we explored the inhibition of rRNA maturation, and found that targeting FBL did not induce any detectable DNA damage. Collectively, these findings demonstrate the potential of targeting the rRNA maturation process as a therapeutic strategy and identify FBL as a promising druggable target.

Ribosome maturation and assembly involves more than 200 factors, many of which are essential and represent additional potential targets, among which we, and others, identified the C/D box snoRNP (13–15). In this study, we show that FBL represents an actionable target. We demonstrated that FBL knock-down prevents the cleavage of the 5’-ETS region in HeLa cells (16), and now in TNBC cellular models. These observations are consistent with the role of FBL in structuring U3 snoRNA, a snoRNA that plays an essential role in the folding and cleavage of the 5’-ETS region of the pre-rRNA (15). It should be noted that FBL is a 2’-O-methyl transferase that modifies about 112 nucleotides in human rRNA (12,19). Future studies will be required to determine whether alterations in 2’-O-methylation contribute to the growth defects observed following FBL knockdown.

Ribosome biogenesis inhibition induces ribosomal stress, a cell response triggered by the accumulation of free ribosomal proteins, mainly uL18 (RPL5) and uL5 (RPL11), and ribosome biogenesis factors such as NPM1 (34), which activate the p53 pathway by trapping MDM2. Nonetheless, while p53 increases the sensitivity of cells to ribosomal stress, it is not required for the ribosomal stress-induced cell growth defect and cell death (35). Accordingly, MDA-MB-231 and BT-20 cells displayed high sensitivity to RNA Pol I inhibitors, with IC_50_ in the low nanomolar range, and to FBL knock-down despite expressing a mutant p53 protein. This represents a key advantage for the development of the therapeutic inhibition of ribosome biogenesis, since the penetrance of *TP53* mutation is over 80% in TNBC (36). In both TNBC cell lines used in this study, cell cycle arrest, but no apoptosis was observed after inhibition of ribosome biogenesis, whether through RNA Pol I inhibition or FBL knockdown. Such a phenotype is not specific to TNBC since apoptosis was not observed after ribosome biogenesis inhibition in several other models of cancer including melanoma (A375 cells), pancreatic cancer (MIA-PaCa cells), osteosarcoma (MNNG and U2OS cells) and prostate cancer (PC3, DU145 and LNCaP cells) (31,35,37). The absence of apoptosis cannot be simply attributed to TP53 mutation. Indeed, there seems to be conflicting data since apoptosis was triggered upon CX-5461 treatment in TP53 mutated osteosarcoma cells lines (31), but was inhibited in lymphoma cells upon TP53 knock-down (38). More globally, the contribution of p53 in the cell response to ribosome biogenesis inhibition remains to be defined (34).

In summary, our findings highlight that targeting ribosome biogenesis represents a promising therapeutic strategy for TNBC, a breast cancer subtype in critical need of novel treatment approaches. Although the development of safer RNA Pol I inhibitors remains a potential avenue, our data strongly support the therapeutic potential of inhibiting ribosome maturation, specifically pre-rRNA cleavage, over rRNA synthesis. Furthermore, we highlight rRNA maturation factors, such as snoRNPs, as viable and druggable targets for anticancer therapy.

## Material and methods

### Cell Culture and shRNA/siRNA Transfection and Treatments

MDA-MB-231 and BT-20 cells were cultured in DMEM-F12 medium (Life Technologies) supplemented with 1% GlutaMAX (Life Technologies), 10% foetal bovine serum (CliniSciences), and 1% penicillin-streptomycin (10,000 U/mL; Life Technologies). Cells were maintained at 37°C in a humidified incubator with 5% CO₂.

MDA-MB-231 and BT-20 cell lines with inducible shRNA expression were generated via lentiviral infection. Lentiviral particles were produced using the vectors pTRIPZ-shRNA-NS and pTRIPZ-shRNA-351067 (FBL-1) and 351068 (FBL-2), acquired from Open Biosystems (Dharmacon). The shRNA sequences used were as follows: shRNA-FBL-1: TGCATCTTTTTCACTTCGG, and shRNA-FBL-2: AGACCATCCGGACCAACGA, and a non-specific shRNA (shRNA-NS) as the control: GCGATCTCGCTTGGGCGAGAGTAAGTA. Post-infection, cell populations were selected with 1 μg/mL puromycin for 14 days. shRNA expression was induced by treating the cells with 1 μg/mL doxycycline. Clonal populations were isolated through single-cell cloning and selected based on FBL expression following shRNA induction.

For siRNA experiments, three siRNA duplexes were used for fibrillarin silencing as previously described (16): “400”:5’- UUU CCU CGA CAA AUG AAG AC – 3’, “882”: 5’- AAA AUC ACA AAG UGU CCU CCA -3’, “1090”: 5’- UCU CUC GCA AUC CUG ACA G -3’. A non-targeting siRNA was used as a negative control (negative control siRNA duplex; Eurogentec SR-CL 000-005). Cells were transfected with these siRNAs using Interferin (Polyplus) following the manufacturer’s instructions. Forty-eight- or seventy-two-hours post-transfection, cells were seeded for subsequent analyses.

CX-5461 was dissolved in 25 mM NaH₂PO₄ (pH 7.4), BMH-21 was dissolved in phospho-citrate buffer (pH 6), as previously described (7,8). Doxycycline was dissolved in water (Life Technologies). Actinomycin D was dissolved in ethanol at 5 mg/mL. All drugs were diluted at working concentration in cell culture medium.

### Cell Viability and IC₅₀ Measurement

IC₅₀ values were determined using real-time growth monitoring with the Incucyte® S3 system (Sartorius®). For CX-5461 and BMH-21 treatments, cells were exposed to the compounds on day 0. For shRNA experiments, shRNA expression was induced with 1 μg/mL doxycycline 2 days prior to monitoring. Cells were seeded onto 96-well plates, and cell confluence was measured every 2 h. The IC₅₀ of RNA polymerase I inhibitors at 96 h was calculated using Prism® software.

### RNA Extraction and RT-qPCR

RNA was extracted using the TRIzol®-chloroform method and quantified with a NanoDrop® 2000 spectrophotometer (Thermo Scientific®). Reverse transcription was performed on 200 ng of purified RNA using the PrimeScript™ RT kit (Takara®, RR037B) with random 6-mer primers. Quantitative PCR (qPCR) was conducted using the LightCycler® 480 SYBR Green I Master Mix (Roche®, 4887352001) with specific primer pairs (see Supplementary Table S1) on the LightCycler® 96 instrument. Relative fold changes in gene expression were calculated using the 2^−ΔΔCT method. The primer sequences were as follows: FBL-Fwd: 5’-CCT GGG GAA TCA GTT TAT GG3’; FBL-Rev: 5’-CCA GGC TCG GTA CTC AAT TTT-3’; GAPDH-Fwd: 5’-GCC CAA TAC GAC CAA ATC C AGC-3’; GAPGH-Rev: 5’-CAC ATC GCT CAG ACA C-3’

### Western blot

Ten to forty micrograms of whole-cell protein extract were separated on 12% or 4%-20% gradient SDS-polyacrylamide gels. Proteins were then transferred onto a 0.2 µm nitrocellulose membrane. The membrane was blocked with 5% nonfat milk in TBST (Tris-buffered saline with 0.1% Tween 20) for 1 h at room temperature. Primary antibodies (listed below) were incubated with the membrane for 1 h at room temperature or overnight at 4°C in 5% milk-TBST. Proteins were detected by chemiluminescence using anti-rabbit or anti-mouse HRP-conjugated secondary antibodies (7074/7076, HRP-linked secondary antibodies, Cell Signaling Technology®) diluted 1:5000, and Clarity™ ECL substrate (Bio-Rad®). Alternatively, proteins were detected using fluorescent secondary antibodies (12004161/12005867, StarBright Blue 700, Bio-Rad®). Images were acquired using the ChemiDoc™ MP Imaging System (Bio-Rad®), and signal intensities were analysed using ImageLab™ software (Bio-Rad®).

### Colony formation assay

Colony-forming assays were performed by seeding 1,000 cells per well in a 6-well plate. After 24 h, cells were treated with different concentrations of RNA polymerase I (Pol I) inhibitors. For shRNA-expressing cells, doxycycline was added at the time of plating. After 10 days, the colonies were fixed with 4% paraformaldehyde for 15 min at room temperature, stained with 1% crystal violet for 2 h at room temperature under gentle agitation, and washed with PBS. Colony counts were performed using ImageJ software with Guzman’s plugin (39).

### Spheroid formation assay

Five hundred cells were plated in 96-well ultra-low attachment (ULA) plates in 2.5% Matrigel medium. For wild-type cells, spheroids were treated with different concentrations of Pol I inhibitors (CX-5461, BMH-21, or Actinomycin D) 24 h post-plating. For shRNA-expressing cells, doxycycline induction was performed 2 days prior to plating. Spheroid growth was monitored using the Opera Phenix® High-Content Screening System (Revvity™). Spheroid size was measured using Signal Image Artist (SImA) software (Revvity™).

### Immunofluorescence (IF)

Cells cultured on 96-well plates or coverslips were fixed with 4% paraformaldehyde for 5 min at room temperature and then permeabilized with 0.5% Triton X-100 for 5 min at room temperature. Cells were blocked with 3% FBS in PBS for 30 min. Primary antibodies (Supplementary table S5) were incubated with the cells for 1 h at room temperature, followed by incubation with fluorescence-labelled secondary antibodies for 30 min. Nuclei were stained with Hoechst (0.5 µg/mL). Images were acquired using an Opera Phenix® High-Content Screening System (Revvity™).

### Cell Cycle Analysis

Cells were cultured in 96-well plates and treated for 18 h with CX-5461, BMH-21, or actinomycin D, or for 3 and 7 days with doxycycline for shRNA induction. Cells were then incubated with 5-Ethynyl-2’-deoxyuridine (EdU, 1 mM) for 1 h at 37°C with 5% CO₂ in a humidified incubator. Following incubation, cells were fixed with 4% paraformaldehyde for 5 min at room temperature and permeabilized with 0.5% Triton X-100 for 20 min at room temperature. Click detection was performed using a fluorescent-azide mix for 30 min, protected from light. Cells were then blocked with 3% FBS in PBS for 30 min and incubated with primary Phospho-Histone H3 (Serine 10) antibody (Merk® #06-570) for 1 h at room temperature. This was followed by incubation with fluorescence-labelled secondary antibodies for 30 min. Nuclei were stained with Hoechst (0.5 µg/mL). Cells were imaged using an Opera Phenix® automated microscope (Revvity™), and cell cycle phases were determined based on EdU signal (S-phase), Phospho-Histone H3 signal (M-phase), and Hoechst staining (G1 and G2/M phases) using SImA software (Revvity™).

### Fluorescence In Situ Hybridization (FISH)

The FISH experiments were performed as described previously (40). MDA-MB-231 and BT-20 cells were seeded on coverslips in 24-well plates. Cells were treated with CX-5461, BMH-20 (1 µM), or Actinomycin D (0.05 µg/mL) for 4 h. Cells were then fixed with 4% formaldehyde for 30 min. Cells were incubated with 2.5 ng of ETS-1 probe for 5 h in a humidified chamber at 37°C. Images were acquired using a Zeiss Axio Imager widefield fluorescence microscope, a Zeiss LSM 980 confocal microscope, and an Opera Phenix® High-Content Screening System (Revvity™). The sequences of the probes were previously published (40). ETS1 probe: AGACGAGAACGCCTGACACGCACGGCAC-AlexaFluor 555. 5’-ITS1 probe: CCTCGCCCTCCGGGCTCCGTTAATGATC-AlexaFluor 488

### Northern Blot

RNA samples were analysed by northern blot as previously described (41). Briefly, total RNAs were resolved by electrophoresis on 1.2% agarose, 6% formaldehyde, 0.02 M 3-(*N*-morpholino) propanesulfonic acid (MOPS) gels. Transfers were performed by capillarity on a nylon membrane (Nytran SuperCharge, Whatman) with 10X SSC (1.5 M sodium chloride, 0.15 M sodium citrate, pH 7.0). Membranes were incubated overnight at 42°C with hybridization buffer (ULTRAhyb™-Oligo, Invitrogen) containing ITS1 or ITS2 DNA probes. Membranes were washed four times with 0.1X SSC and 0.1% SDS before signal exposure on ChemiDoc MP (Bio-Rad). Quantification was performed using Image Lab software (Bio-Rad) and relative 47S pre-rRNA levels were determined by normalisation against intermediate ARNr levels. The probe sequences were as follows:

ITS1 5′- CCGCGGGCCTCGCCCTCCGGGCTCCGTTAATGATCCTTCC-Dylight 800.
ITS2 5′- CTGCGAGGGAACCCCCAGCCGCGCA-Dylight 800

### Cell Death detection with DRAQ7

Two thousand cells (MDA-MB-231 and BT-20) per condition were cultured in 96-well plates. Cells were treated with RNA Pol I inhibitors one day after plating for 3 days, or at the time of plating for shRNA-expressing cells for 4 days. Subsequently, cells were stained with DRAQ7 (1:1000, 424001 BioLegend® Inc.) and nuclei were counterstained with Hoechst (0.5 µg/mL) for 3 h. The cells were then imaged using an Opera Phenix® High-Content Screening System (Revvity™), and DRAQ7(positive cells were quantified using SImA software (Revvity™).

### Caspase 3/7 Incucyte® Assay

Two thousand cells (MDA-MB-231 and BT-20) per condition were plated in 96-well plates. Cells were treated 24 h post-plating with Pol I inhibitors (CX-5461, BMH-21, actinomycin D), or directly with doxycycline for shRNA-expressing cells, including a 2-day pre-treatment. Incucyte® Caspase 3/7 Green Dye was added to the medium as recommended by the manufacturer (Thermo Fisher Scientific®). Cells were monitored in real-time using the Incucyte® S3 system (Sartorius®) for one week, with images captured every 2 h. Apoptotic cells were quantified using Incucyte® software.

### Xenograft Models

Five-week-old female athymic nude mice were obtained from JANVIER® and maintained under specific pathogen-free conditions. The animal protocol described below was reviewed and approved by the Animal Ethical and Welfare Committee (protocol number: 2021101416573294). A total of 2 × 10^6^ MDA-MB-231 cells or 3 × 10^6^ BT-20 cells (normal [WT or shNC] or FBL-depleted [shFBL-1, -2]) were inoculated subcutaneously into the dorsal flank of the nude mice in PBS. Treatment with CX-5461, BMH-21, doxycycline, or vehicle was initiated once the tumors reached a similar size (approximately 100 mm³). Mice were randomly assigned to control and treatment groups. CX-5461 (50 mg/kg) dissolved in 25 mM NaH₂PO₄ (pH 7.4) was administered via oral gavage three times per week. BMH-21 (50 mg/kg) dissolved in phospho-citrate (pH 6.0) was administered via intraperitoneal injection five times per week. Doxycycline (2 mg/mL) dissolved in water with 3% sucrose was provided in drinking water continuously. Mouse weight was monitored weekly, and tumor size was measured three times per week using callipers. Animal experimental procedures and immunohistochemical staining were carried out by the Small Animal Platform (P-PAC) and the Pathology-Research Platform of the Cancer Research Center of Lyon (CRCL).

### Trypsin digestion and mass spectrometry analysis

For mass spectrometry analysis, 5 independent whole cell extracts were prepared from MDA-MB-231 cells untreated or treated for 18 h with CX-5461 at 700 nM, BMH-21 at 1000nM, or Actinomycin D at 0.05µg/mL, using a lysis buffer containing 62.5 mM Tris-HCl, 1% SDS and 1 mM DTT. Protein concentration was measured using TCA precipitation assay. Between 40 µg and 100µg of proteins were subjected to a disulfide bridge reduction for 10 min at 95 °C under agitation followed by an alkylation of cysteine residues in 60 mM iodoacetamide for 30 min in the dark at room temperature. Each reduced/alkylated protein sample was then digested using the S-Trap^TM^ micro spin column protocol [10.1021/acs.jproteome.8b00505]. Briefly, equivalent volumes of 10% SDS were added to each sample in order to reach a final SDS concentration of 5%. Undissolved matter was removed by centrifugation for 8 min at 13,000× g. Aqueous phosphoric acid (12%) was added at a ratio of 1:10 to the protein sample for a final concentration of ∼1.2% phosphoric acid, followed by seven volumes of S-Trap binding buffer (90% methanol, 100 mM TEAB, pH 7.1). After gentle mixing, the protein solution was loaded into an S-Trap filter several times, each separated by a 4000×g centrifugation step, until all the SDS lysate/S-Trap buffer had passed through the column. Afterwards, the captured proteins were washed six times with 150μL S-Trap binding buffer. Digestion was performed overnight at 37 °C by the addition of 20μL of trypsin (Sequencing Grade Modified Trypsin, Promega) at 37.5ng/μL in 50 mM ammonium bicarbonate. The digested peptides were eluted by the addition of 35μL of 50 mM ammonium bicarbonate and 1 min centrifugation at 4000×g, followed by 35μL of 0.2% formic acid and 4000×g centrifugation (1 min) and finally 40μL of 50% aqueous acetonitrile containing 0.2% formic acid and a last 1 min centrifugation step at 4000×g. Before drying down the pooled eluates under speed vacuum, the peptide amount was measured using the Pierce Quantitative Fluorometric Peptide assay (Thermo Fisher Scientific®).

Based on the fluorometric peptide assay, samples were resuspended in 2% acetonitrile (ACN), 0.2% formic acid (FA) at a concentration of 250 ng/μL and sonicated for 10 min before analysis by online nanoLC-MS/MS using an UltiMate® 3000 RSLCnano LC system (Thermo Scientific, Dionex) coupled to an Orbitrap Exploris 480 mass spectrometer using FAIMS Pro Duo interface (Thermo Fisher Scientific, Bremen, Germany). One microliter of each sample was directly loaded onto the analytical C18 nanocolumn (Acclaim PepMap RSLC C18 75 µm ID x 50 cm, 2µm, 100Å, Thermo Fisher Scientific) heated at 45 °C and equilibrated in 97.5% solvent A (0.2% formic acid) and 2.5% solvent B (80% ACN, 0.2% formic acid) during 16 min at a flow rate of 300 nL/min. Peptides were eluted using a gradient of 2.5% to 25% of solvent B for 102 min, then 25% to 40% of solvent B for 20 min at a flow rate of 300 nL/min (165 min running time and 122 min total gradient time). The Orbitrap Exploris 480 was operated in FAIMS mode (Standard Resolution, gas flow of 3.9 L/min and two compensation voltages used: −45V and −60V) and in data-dependent acquisition mode using a top-speed approach (cycle times of 0.8s and 0.7s, respectively for each CV) with the Xcalibur software. For both CV experiments, the following parameters were used: Survey scans MS were acquired in the Orbitrap on the 375 – 1200 m/z range, with the resolution set to a value of 60 000, a « Normalized AGC target » at 300% and a « Maximum injection time » at 60ms for −45V and 50ms for −60V. The most intense multiply charged ions (2+ to 6+) above a threshold ion count of 5 × 103 were selected, for fragmentation by higher energy collisional dissociation with an isolation window of 1.6 m/z and a normalized collision energy of 30%. The resulting fragments were analysed in the Orbitrap at 15,000 resolution, a « Normalized AGC target » at 100% and a « Maximum injection time » set to Auto. Dynamic exclusion was used within 45 s with a 10ppm tolerance, to prevent repetitive selection of the same peptide. For internal calibration the 445.120025 ion was used as lock mass in the first MS Experiment for the first five minutes using a CV at −30V with a 120,000 resolution.

### MS-Based Protein Identification and Label-Free Quantitative Proteomics Analysis

All raw MS files were processed with MaxQuant software (v 2.5.1.0) for database search with the Andromeda search engine and for quantitative analysis. Spectra were searched against the human proteins in the Uniprot Reference Proteome database (database release version of April 2023), containing 81,791 sequences (downloaded from http://www.uniprot.org), and the set of common contaminants provided by MaxQuant. Carbamidomethylation of cysteines was set as a fixed modification, whereas oxidation of methionine and protein N-terminal acetylation were set as variable modifications. Specificity of trypsin digestion was set for cleavage after K or R, and two missed trypsin cleavage sites were allowed. The precursor mass tolerance was set to 20ppm for the first search and 4.5 ppm for the main Andromeda database search. The mass tolerance in tandem MS mode was set to 20 ppm. The minimum peptide length was set to seven amino acids, and the minimum number of unique or razor peptides was set to one for validation. Andromeda results were validated by the target decoy approach using a reverse database, with a false discovery rate set at 1% at both PSM (peptide sequence match) and protein level. For label-free relative quantification of the samples, the match between runs option of MaxQuant was enabled with a match time window of 0.7min, to allow cross-assignment of MS features detected in the different runs, after alignment of the runs with a time window of 20 min. Proteins were quantified by the MaxLFQ algorithm integrated in the MaxQuant software. A minimum ratio count of two unique or razor peptides was required for quantification.

Statistical analysis of the proteomic data was performed in the Perseus software (version 1.6.15.0) after loading the “proteinGroups.txt” files from MaxQuant. Reverse database hits were removed as well as proteins identified as contaminants. LFQ intensities were log2 transformed and replicate samples were grouped. Proteins with less than three valid values in at least one group were removed and missing values were imputed from a normal distribution around the detection. For each pairwise comparison, an unpaired two-tailed Student t test was performed, and proteins were considered differentially abundant when their absolute log2-transformed fold change (FC) was higher than 1 and their p-value lower than 0.05. Volcano plots were drawn to visualize significant protein abundance variations between the compared conditions. These represent −log_10_ (p-value) according to the log_2_ ratio. The complete list of the identified and quantified proteins and analysed according to this statistical procedure is described in Table S6.

### TCGA Breast Cancer cohorts and statistical analyses

Gene expression BRCA dataset were uploaded from the TCGA database (25). Gene expression was given as “log2(norm_count + 1)”. After data upload and transformation, a BRCA tumor dataset was built by conserving only tumoral tissues collected at diagnosis from women without any neoadjuvant treatment at primary site and for which status of either PAM50 or oestrogen, progesterone and HER2 receptors was available. The gene signatures were derived by calculating the median expression of genes involved in each signature per tumor, the RiBi signature being composed of 240 genes (Supplementary Table 1 “signature”) and the snoRNP signature of 4 genes. For each gene signature, mean comparison was performed between four breast cancer subtypes. Two classifications were used: PAM50 as given by the TCGA database; and intrinsic breast cancer status determined using the oestrogen, progesterone and HER2 receptors’ IHC data given by the TCGA database (Supplementary Tables S2-S3). Prior to mean comparison, application conditions of statistical tests were performed (normality: Shapiro-Wilk test; homogeneity of variances: Levene test). Comparison between all breast cancer subtypes was performed using ANOVA. If significant, a two-by-two comparison was assessed using either parametric Student t-tests or nonparametric Mann-Whitney tests depending on results of the control of the application conditions. P-values of two-by-two comparison were then adjusted to multiple testing using False Discovery Rate method. Statistical significance was based on a p < 0.05 where the H0 hypothesis is rejected. Violin plots were used to visualize the distribution per breast cancer subtype. Statistical analyses and visualisations were performed using R Studio (version 2024.04.1+748).

### Statistical analysis and graphical representations

Statistical analyses (ANOVA, Student t-test) and graphical representations were performed with GraphPad PRISM 10.

## Acknowledgements

We thank the core facilities of the Cancer Research Center of Lyon for technical help: in particular Mélina GAUTIER (PIC Cellular Imaging Platform of CRCL), Isabelle GODDARD (P-PAC animal core facility), Nicolas GADOT (PAR research pathology platform) and Maxime GARCIA (Center for Drug Discovery and Development, Centre Léon Bérard). We thank Brigitte Manship for editing the manuscript. We are grateful to Thierry DUBOIS (Institut Curie, Paris, France) for providing the TNBC cell lines. The study was supported by grants from: La ligue régionale Contre le Cancer région AURA to FC, ANR (Actimeth, 19-CE12-0004; LabEx DEVweCAN, ANR-10-LABX-0061; Institut Convergence ANR-17-CONV-0002), European network COST Translacore (COST Action CA21154). This work was supported in part by the French Ministry of Research with the Investissement d’Avenir Infrastructures Nationales en Biologie et Santé program (ProFi, Proteomics French Infrastructure project; ANR-10-INBS-08). C.J. and J.R. received a PhD fellowship from the Ligue Nationale Contre le Cancer. P.L.M. received a PhD fellowship from the French Ministry for higher education and research. FNVL received a PhD fellowship from Ligue Nationale Contre le Cancer and Fondation pour la Recherche Médicale. F.C. is a CNRS research fellow. V.M., S.D. and J.-J.D. are Inserm research fellows.

## Authors contributions

CJ, PLM, AG, CI, FB, M-AM, MM, MCD, CF, JM, LB and SD performed experiment and analysed data. FNVL, TF, VM and FC analysed data. CV contributed to methodology. FC, CJ, SG and VM Conceived the study. FC, JJD and VM secured funding. JJD and VM reviewed the manuscript.

CJ, PLM and FC wrote and reviewed the manuscript.

## Data and material availability

The mass spectrometry proteomics data have been deposited to the ProteomeXchange Consortium via the PRIDE(42) partner repository with the dataset identifier PXD060027. All other raw data are available upon request from the corresponding author.

Biological material is available upon reasonable demand.

## Ethics statement

The animal protocol described in this study was reviewed and approved by the Animal Ethical and Welfare Committee (protocol number: 2021101416573294).

