## Supplementary Figures for "Fibrillarin-mediated ribosomal RNA maturation is a novel therapeutic vulnerability in triple-negative breast cancer"

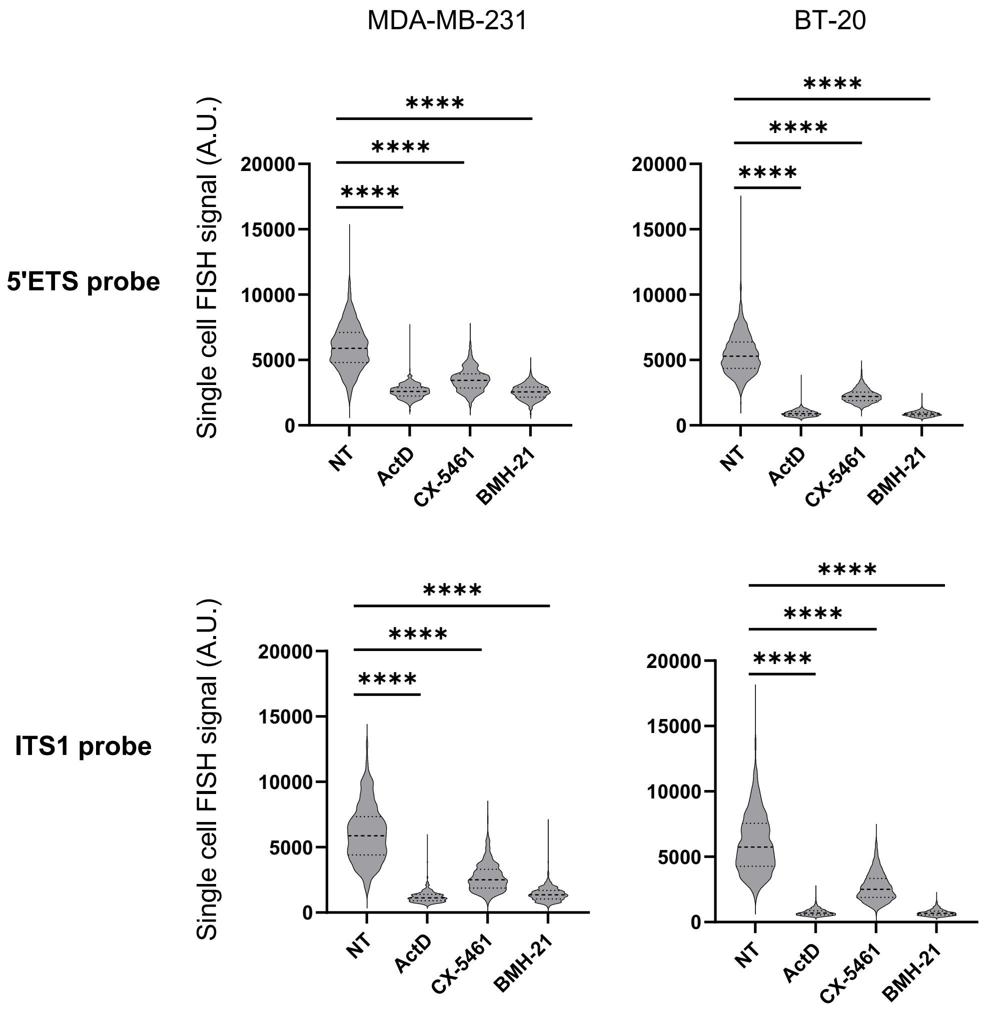


**Supplementary Figure 1, related to Figure 2**

Violin plot showing the quantification of FISH signal in single cells, from figure 2A. FISH was performed using probes specific for 5'ETS and ITS1 pre-rRNA regions on MDA-MB-231 and BT-20 cells either untreated (NT) or treated with 1 μM CX-5461, 1 μM BMH-21, or 0.05 μg/mL actinomycin D for 4 h. Welch's ANOVA test. $p < 0.001$ ****

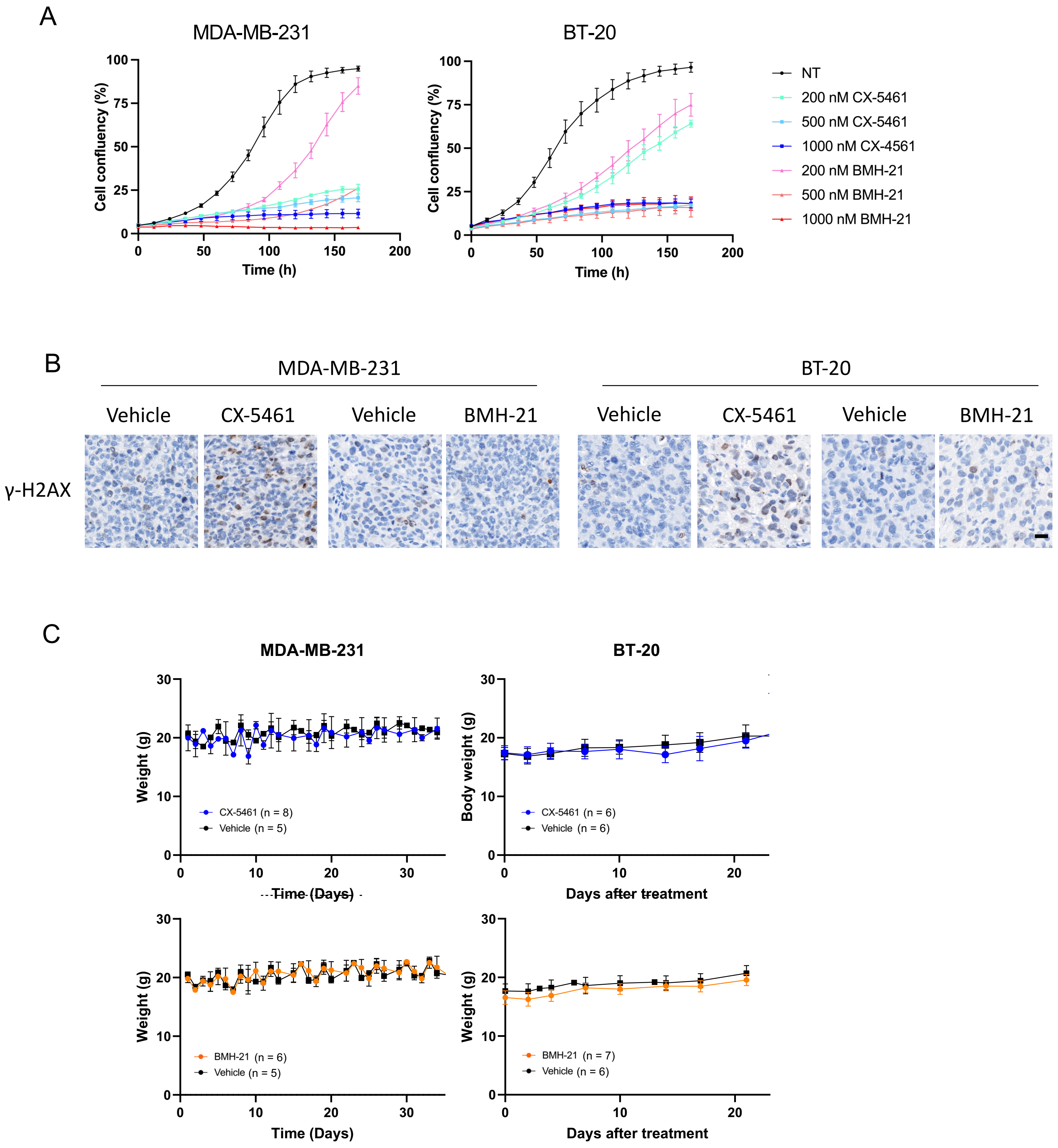


**Supplementary Figure 2, related to Figure 3**

**A)** Representative proliferation curves obtained by real-time imaging, following treatment with 1 nM to 1000 nM of CX-5461 or BMH-21. Each curve is representative of at least two independent biological replicates. Data are mean values +/- s.d.

**B)** Representative images of immunohistochemical staining for γ-H2AX in MDA-MB-231 and BT-20 xenografts. Xenografts are from the experiment described in Figure 2F and Figure 3B. Scale bar: 20 µm.

**C)** Weight of mice untreated (vehicle) or treated with BMH-21 or CX-5461, presented in Figure 3F. Data are mean values +/- s.e.m.

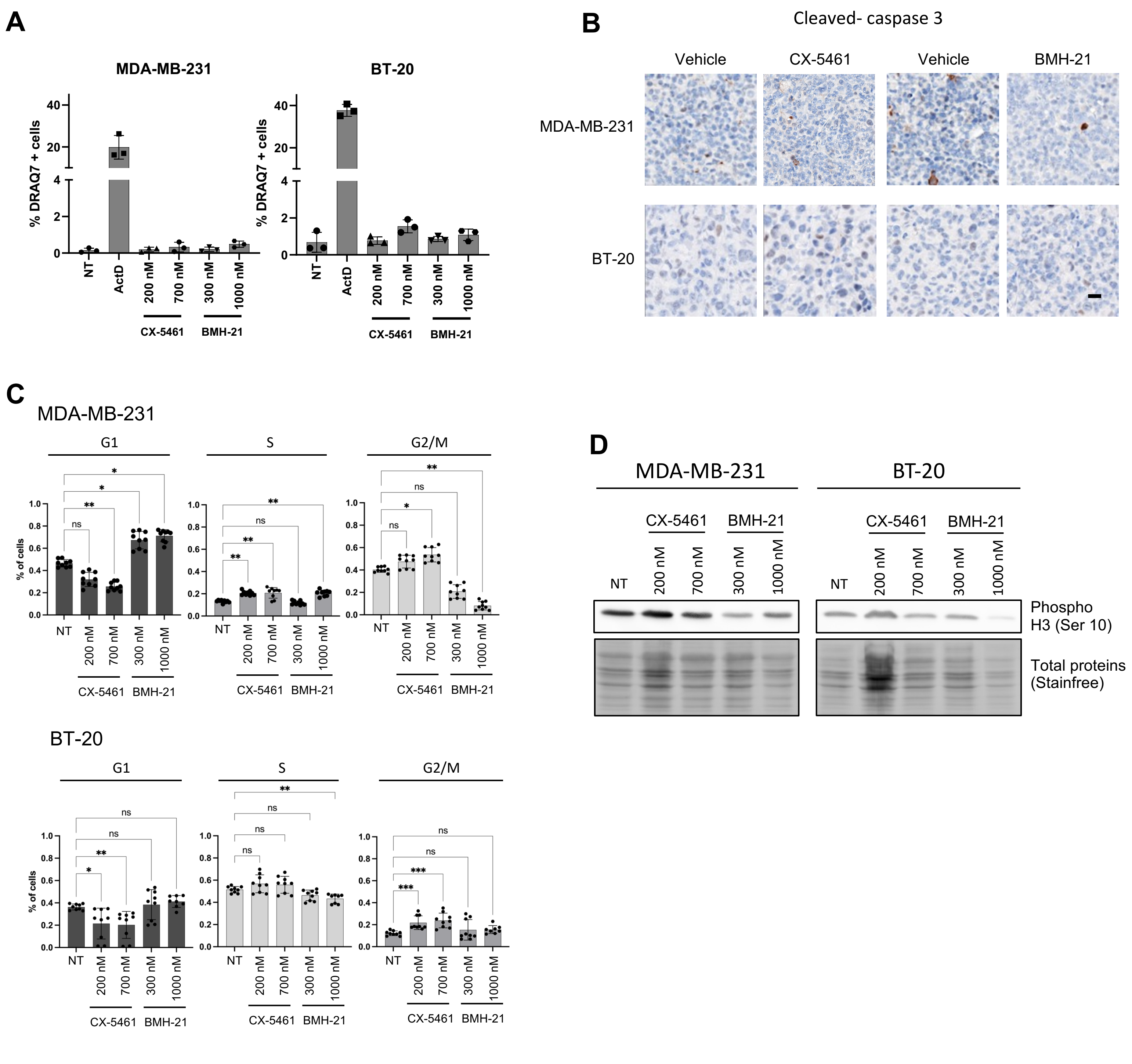


**Supplementary Figure 3, related to Figure 4A-B**

**A)** Quantification of dead cells (DRAQ7 positive cells) using the DRAQ7 fluorescent dye, on MDA-MB-231 and BT-20 cells, either untreated or treated with CX-5461 or BMH-21 at the indicated concentration for 72h. Data are mean values +/- sd from 3 independent replicates.

**B)** IHC staining of cleaved caspase 3, on sections of xenograft tumors, from the experiment shown in Figure 3F. Images are representative of 5 biological replicates. Scale bar: 50 µm

**C)** Proportion of cells in G1, S, and G2 phases of the cell cycle after overnight treatment (18 h) in MDA-MB-231 and BT-20 cell lines untreated (NT) or treated 18h with either CX-5461 or BMH-21 at the indicated concentration. Statistics were performed on the 3 technical replicates of the 3 biological replicates. Kruskal-Wallis test. * $p < 0.05$, ** $p < 0.01$, *** $p < 0.001$ **** $p < 0.0001$.

**D)** Immunoblot of phospho-Histone H3 (Ser10) in MDA-MB-231 and BT-20 cell lines either untreated (NT) or treated 72h with CX-5461, or BMH-21, at the indicated concentration. Data representative of 2 independent biological replicates.

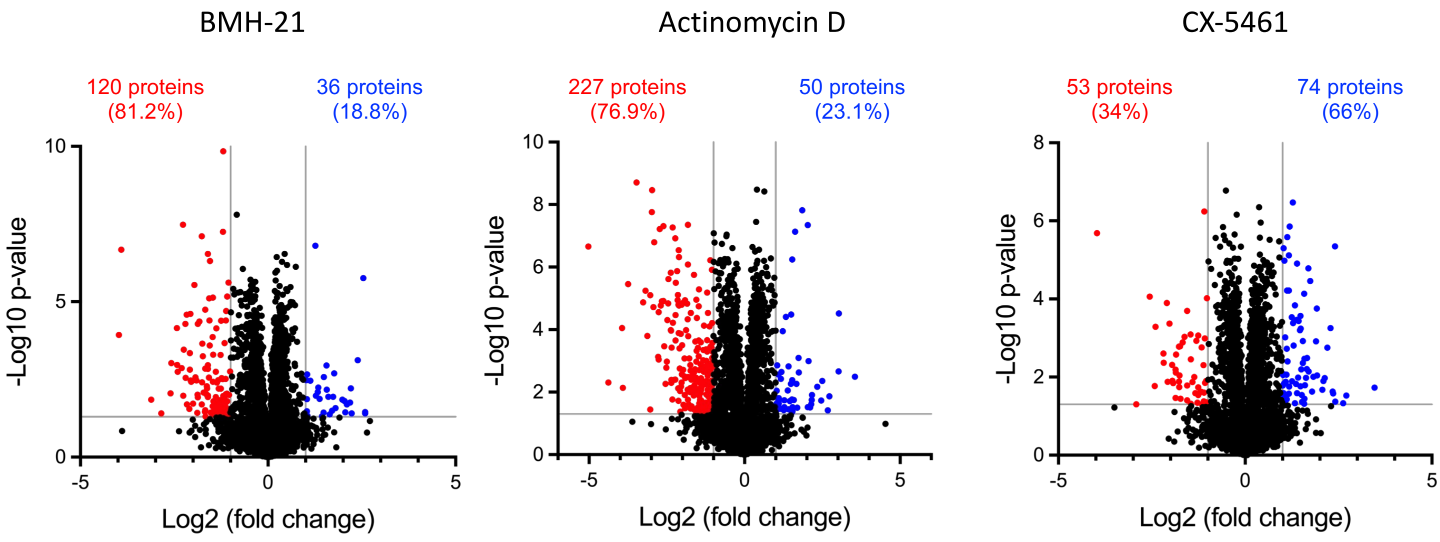


**Supplementary figure 4, related to figure 4C**

Volcano plot of protein levels observed in the proteomic analysis of MDA-MB-231 cells treated with CX-5461, BMH-21 and actinomycin D, presented in Figure 4C. Significantly changed level of proteins are colored (blue, up-regulated, red, down-regulated) based on a log2 fold change of more than 1 and a p-value lower than 0.05 (Log10 > 1.3; Student t-test, n = 5).

The number of significantly different proteins are indicated on top, with their respective percentage).

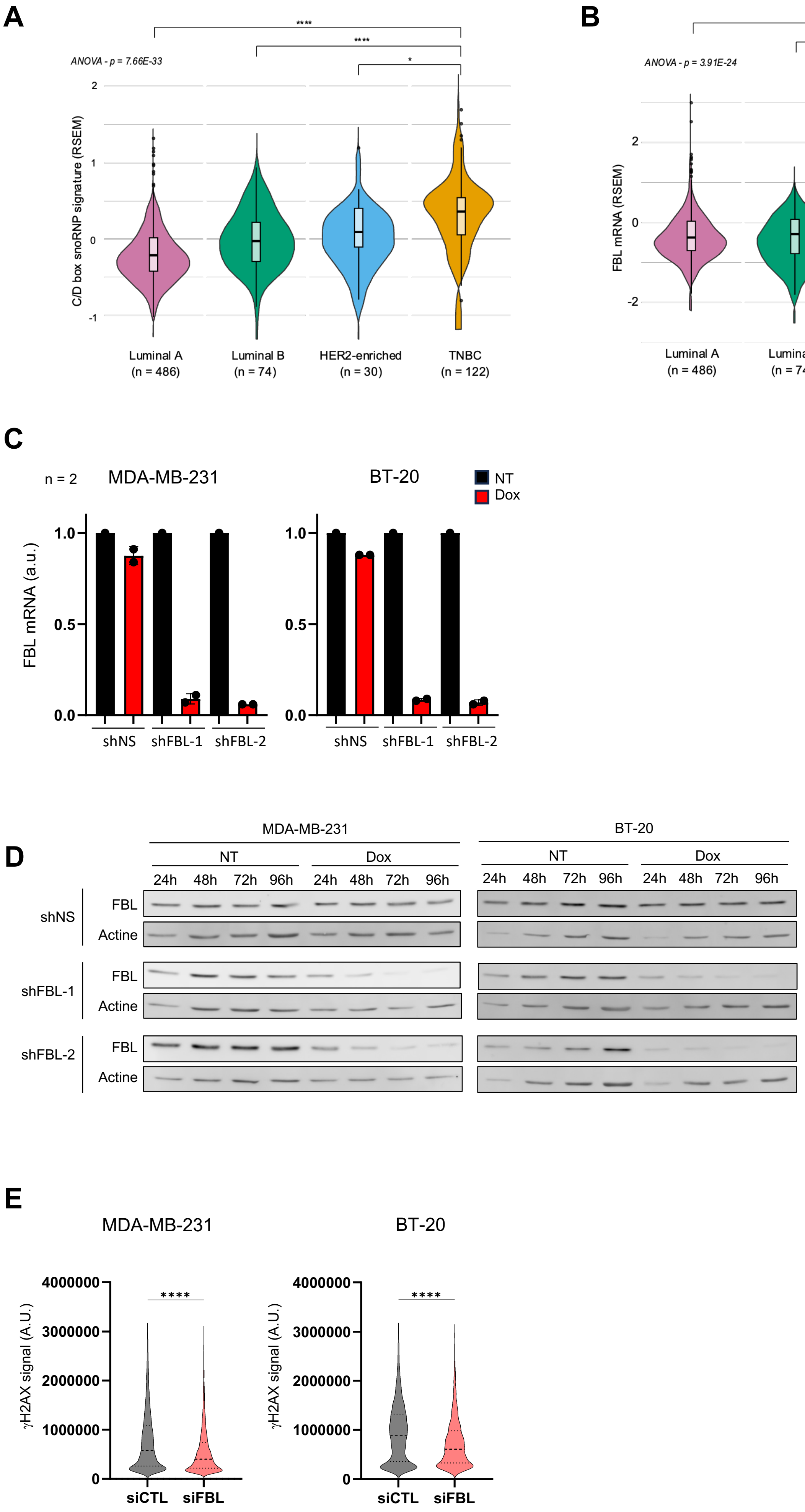

A
ANOVA - p = 7.66E-33
C/D box snoRNP signature (RSEM)
Luminal A
(n = 486)
Luminal B
(n = 74)
HER2-enriched
(n = 30)
TNBC
(n = 122)
B
ANOVA - p = 3.91E-24
FBL mRNA (RSEM)
Luminal A
(n = 486)
Luminal B
(n = 74)
HER2-enriched
(n = 30)
TNBC
(n = 122)
C
n = 2
MDA-MB-231
BT-20
NT
Dox
FBL mRNA (a.u.)
shNS
shFBL-1
shFBL-2
D
MDA-MB-231
BT-20
NT
Dox
24h
48h
72h
96h
shNS
shFBL-1
shFBL-2
FBL
Actine
E
MDA-MB-231
BT-20
γH2AX signal (A.U.)
siCTL
siFBL

**Supplementary figure 5, related to figure 5**

**A-B,** Expression of the C/D box snoRNP protein genes (**A**) and FBL (**B**) mRNA expression levels among the different breast cancer subtypes. Normalized transcriptomic data were extracted from the UCSC XENA database. The breast cancer subtypes were categorized using the histological classification, as in figure 1B. n: number of patients; **** $p < 0.0001$ (Anova test, Mann-Whitney test).

**C,** RT-qPCR analysis of FBL mRNA expression in MDA-MB-231 and BT-20 cell lines after 96-h doxycycline-mediated shRNA induction. Cell lines expressing either one of the shRNA targeting FBL (FBL-1 and FBL-2) either a non-targeting shRNA (shNS). Data are mean of two independent biological replicates.

**D,** Western blot analysis of FBL protein levels in MDA-MB-231 and BT-20 cell lines after 24-, 48-, 72- or 96-hour doxycycline-mediated shRNA induction. Cell lines expressing either one of the shRNA targeting FBL (FBL-1 and FBL-2) either a non-targeting shRNA (shNS).

**E,** Single cell immunofluorescence signal of $\gamma$H2Ax, extracted from the experiment shon in Figure 5G.

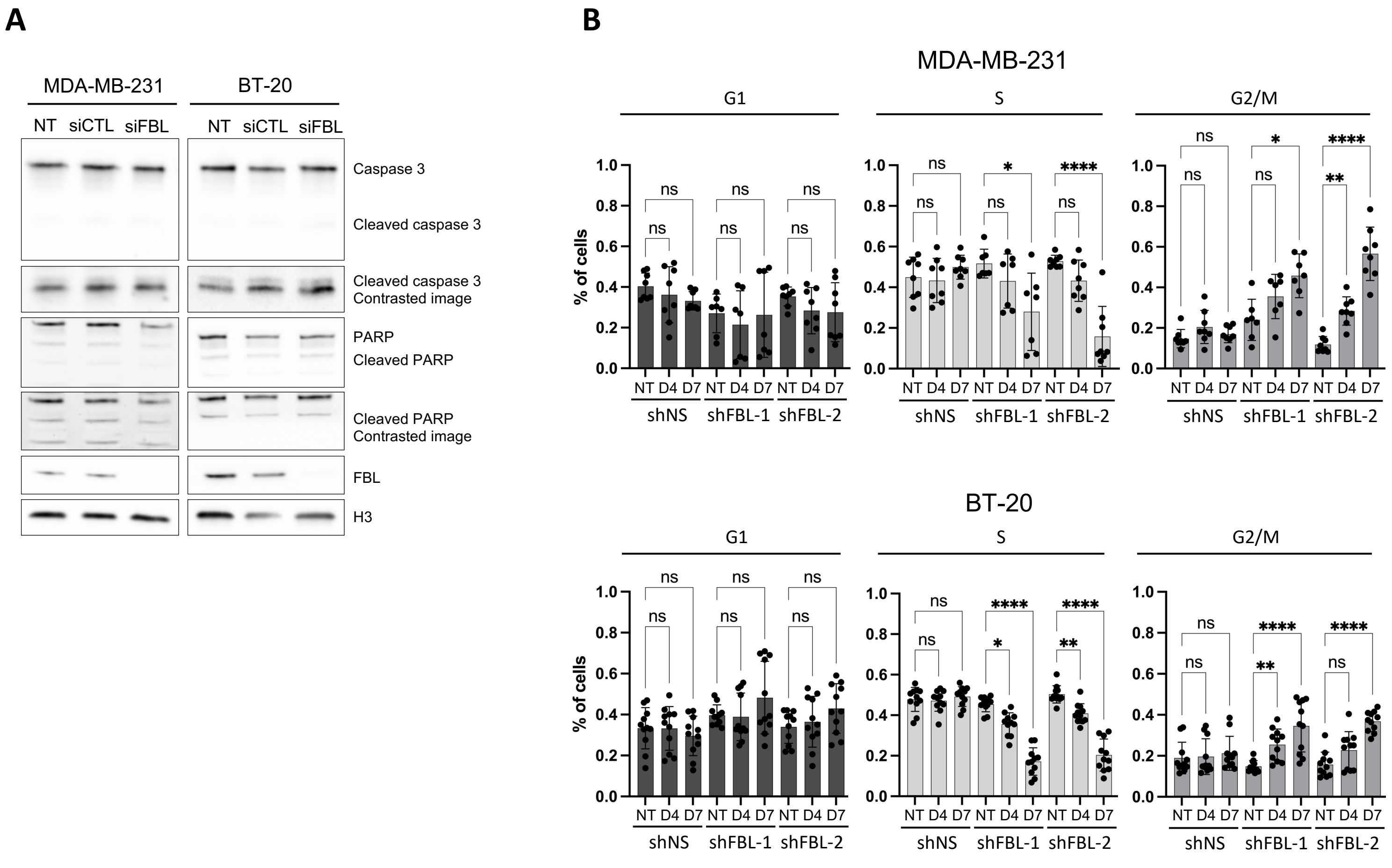


**Supplementary Figure 6, related to Figure 7**

**A,** Western blot analysis of apoptosis-related protein Caspase 3 and PARP in MDA-MB-231 and BT-20 cell lines following doxycycline-mediated siRNA induction for 72 h. The full length and cleaved forms are indicated. Contrasted images are also shown for the cleaved forms. FBL-siRNA efficiency is revealed by FBL detection. Histone H3 is shown as loading control.

**B,** Cell cycle analysis data of the three biological replicates related to the experiment shown in Figure 7C. Data are mean +/- s.d. and individual data point of the proportion of cells in G1, S, and G2/M phases of the cell cycle in MDA-MB-231 and BT-20 cell lines either untreated (NT) or following doxycycline-mediated shRNA induction for 4 (D4) or 7 (D7) days. Statistics were performed on the 3 technical replicates of the 3 biological replicates using a Kruskal-Wallis test. * $p < 0.05$, ** $p < 0.01$, *** $p < 0.001$, **** $p < 0.0001$.

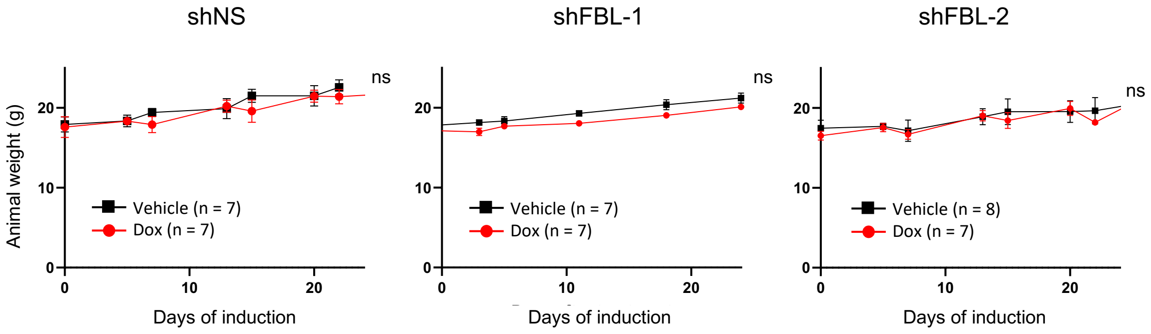


**Supplementary Figure 7, related to Figure 8**
Weight of mice either untreated (vehicle) or treated with doxycycline in the dreaking water, presented in Figure 8A. Data are mean values +/- s.e.m.
