## Supplementary Tables for "Fibrillarin-mediated ribosomal RNA maturation is a novel therapeutic vulnerability in triple-negative breast cancer"

**Supplementary tables S1 to S5**

**Supplementary Table S1**

"Ribosome biogenesis in eukaryotes" gene signature, defined by the KEGG pathway.

| Symbol | Name |
|---|---|
| AATF | apoptosis antagonizing transcription factor |
| ABCE1 | ATP-binding cassette, sub-family E (OABP), member 1 |
| ABCF2 | ATP-binding cassette, sub-family F (GCN20), member 2 |
| ABT1 | activator of basal transcription 1 |
| ANKZF1 | ankyrin repeat and zinc finger domain containing 1 |
| DDX47 | DEAD (Asp-Glu-Ala-Asp) box polypeptide 47 |
| ATP8A1 | ATPase, aminophospholipid transporter (APLT), class I, type 8A, member 1 |
| ATP8A2 | ATPase, aminophospholipid transporter, class I, type 8A, member 2 |
| BCCIP | BRCA2 and CDKN1A interacting protein |
| BMS1 | BMS1 homolog, ribosome assembly protein (yeast) |
| BRIX1 | BRX1, biogenesis of ribosomes, homolog (S. cerevisiae) |
| BYSL | bystin-like |
| C12orf45 | chromosome 12 open reading frame 45 |
| C14orf169 | chromosome 14 open reading frame 169 |
| C1orf109 | chromosome 1 open reading frame 109 |
| EMG1 | EMG1 nucleolar protein homolog (S. cerevisiae) |
| CASC1 | cancer susceptibility candidate 1 |
| EIF4A1 | eukaryotic translation initiation factor 4A1 |
| CEBPZ | CCAAT/enhancer binding protein (C/EBP), zeta |
| IPO4 | importin 4 |
| CINP | cyclin-dependent kinase 2 interacting protein |
| CIRH1A | cirrhosis, autosomal recessive 1A (cirhin) |
| CMSS1 | cms1 ribosomal small subunit homolog |
| COIL | coilin |
| CRBN | cereblon |
| CSNK2A2 | casein kinase 2, alpha prime polypeptide |
| CSNK2B | casein kinase 2, beta polypeptide |
| DBR1 | debranching enzyme homolog 1 (S. cerevisiae) |
| DCAF13 | DDB1 and CUL4 associated factor 13 |
| DDX10 | DEAD (Asp-Glu-Ala-Asp) box polypeptide 10 |
| DDX18 | DEAD (Asp-Glu-Ala-Asp) box polypeptide 18 |
| DDX21 | DEAD (Asp-Glu-Ala-Asp) box helicase 21 |
| DDX24 | DEAD (Asp-Glu-Ala-Asp) box polypeptide 24 |
| DDX27 | DEAD (Asp-Glu-Ala-Asp) box polypeptide 27 |
| DDX28 | DEAD (Asp-Glu-Ala-Asp) box polypeptide 28 |
| DDX31 | DEAD (Asp-Glu-Ala-Asp) box polypeptide 31 |
| DDX49 | DEAD (Asp-Glu-Ala-Asp) box polypeptide 49 |
| DDX5 | DEAD (Asp-Glu-Ala-Asp) box helicase 5 |
| DDX51 | DEAD (Asp-Glu-Ala-Asp) box polypeptide 51 |
| DDX52 | DEAD (Asp-Glu-Ala-Asp) box polypeptide 52 |
| DDX54 | DEAD (Asp-Glu-Ala-Asp) box polypeptide 54 |
| DDX55 | DEAD (Asp-Glu-Ala-Asp) box polypeptide 55 |
| DDX56 | DEAD (Asp-Glu-Ala-Asp) box helicase 56 |
| DHX15 | DEAH (Asp-Glu-Ala-His) box polypeptide 15 |
| DHX33 | DEAH (Asp-Glu-Ala-His) box polypeptide 33 |
| DHX37 | DEAH (Asp-Glu-Ala-His) box polypeptide 37 |
| DIEXF | digestive organ expansion factor homolog (zebrafish) |
| DIP2A | DIP2 disco-interacting protein 2 homolog A |
| DKC1 | dyskeratosis congenita 1, dyskerin |
| DNAJC21 | DnaJ (Hsp40) homolog, subfamily C, member 21 |
| DROSHA | drosha, ribonuclease type III |
| DUSP12 | dual specificity phosphatase 12 |
| EBNA1BP2 | EBNA1 binding protein 2 |
| EFTUD1 | elongation factor Tu GTP binding domain containing 1 |
| EIF3J | eukaryotic translation initiation factor 3, subunit J |
| EIF4A3 | eukaryotic translation initiation factor 4A3 |
| EIF6 | eukaryotic translation initiation factor 6 |
| ERAL1 | Era G-protein-like 1 (E. coli) |
| EXOSC10 | exosome component 10 |
| FBL | fibrillarin |
| FCF1 | FCF1 small subunit (SSU) processome component homolog (S. cerevisiae) |
| FPR3 | formyl peptide receptor 3 |
| FTSJ2 | FtsJ RNA methyltransferase homolog 2 (E. coli) |
| GAR1 | GAR1 ribonucleoprotein homolog (yeast) |
| GNL2 | guanine nucleotide binding protein-like 2 (nucleolar) |
| GNL3 | guanine nucleotide binding protein-like 3 (nucleolar) |
| GRWD1 | glutamate-rich WD repeat containing 1 |
| GTPBP4 | GTP binding protein 4 |
| GTPBP5 | GTP binding protein 5 (putative) |
| HEATR1 | HEAT repeat containing 1 |
| HSP90AA1 | heat shock protein 90kDa alpha (cytosolic), class A member 1 |
| HSPA1A | heat shock 70kDa protein 1A |
| HSPA2 | heat shock 70kDa protein 2 |
| HSPA6 | heat shock 70kDa protein 6 (HSP70B') |
| HSPA8 | heat shock 70kDa protein 8 |
| IGHMBP2 | immunoglobulin mu binding protein 2 |
| IMP3 | IMP3, U3 small nucleolar ribonucleoprotein, homolog (yeast) |
| IMP4 | IMP4, U3 small nucleolar ribonucleoprotein, homolog (yeast) |
| IPO5 | importin 5 |
| IPO7 | importin 7 |
| IPO9 | importin 9 |
| ISG20 | interferon stimulated exonuclease gene 20kDa |
| ISG20L2 | interferon stimulated exonuclease gene 20kDa-like 2 |
| KDM8 | lysine (K)-specific demethylase 8 |
| KIAA0020 | KIAA0020 |
| KPNB1 | karyopherin (importin) beta 1 |
| KRI1 | KRI1 homolog (S. cerevisiae) |
| KRR1 | KRR1, small subunit (SSU) processome component, homolog (yeast) |
| LSG1 | large subunit GTPase 1 homolog (S. cerevisiae) |
| LTN1 | listerin E3 ubiquitin protein ligase 1 |
| LTV1 | LTV1 homolog (S. cerevisiae) |
| MAK16 | MAK16 homolog (S. cerevisiae) |
| MDN1 | MDN1, midasin homolog (yeast) |
| METTL18 | methyltransferase like 18 |
| METTL5 | methyltransferase like 5 |
| MINA | MYC induced nuclear antigen |
| MPHOSPH10 | M-phase phosphoprotein 10 (U3 small nucleolar ribonucleoprotein) |
| MRM1 | mitochondrial rRNA methyltransferase 1 homolog (S. cerevisiae) |
| MRTO4 | mRNA turnover 4 homolog (S. cerevisiae) |
| MTG1 | mitochondrial GTPase 1 homolog (S. cerevisiae) |
| NAF1 | nuclear assembly factor 1 homolog (S. cerevisiae) |
| NAT10 | N-acetyltransferase 10 (GCN5-related) |
| NCL | nucleolin |
| NEMF | nuclear export mediator factor |
| NGDN | neuroguidin, EIF4E binding protein |
| NHP2 | NHP2 ribonucleoprotein |
| NHP2L1 | NHP2 non-histone chromosome protein 2-like 1 (S. cerevisiae) |
| NIP7 | nuclear import 7 homolog (S. cerevisiae) |
| NMD3 | NMD3 homolog (S. cerevisiae) |
| NOA1 | nitric oxide associated 1 |
| NOB1 | NIN1/RPN12 binding protein 1 homolog (S. cerevisiae) |
| NOC4L | nucleolar complex associated 4 homolog (S. cerevisiae) |
| NOL12 | nucleolar protein 12 |
| NOL6 | nucleolar protein family 6 (RNA-associated) |
| NOL8 | nucleolar protein 8 |
| NOL9 | nucleolar protein 9 |
| NOLC1 | nucleolar and coiled-body phosphoprotein 1 |
| NOP10 | NOP10 ribonucleoprotein |
| NOP14 | NOP14 nucleolar protein |
| NOP16 | NOP16 nucleolar protein |
| NOP2 | NOP2 nucleolar protein |

| | |
|---|---|
| NOP56 | NOP56 ribonucleoprotein |
| NOP58 | NOP58 ribonucleoprotein |
| NOP9 | NOP9 nucleolar protein |
| NPAS4 | neuronal PAS domain protein 4 |
| NPLOC4 | nuclear protein localization 4 homolog (S. cerevisiae) |
| NSA2 | NSA2 ribosome biogenesis homolog (S. cerevisiae) |
| NSUN4 | NOP2/Sun domain family, member 4 |
| NSUN5 | NOP2/Sun domain family, member 5 |
| NUFIP1 | nuclear fragile X mental retardation protein interacting protein 1 |
| NVL | nuclear VCP-like |
| NXT1 | NTF2-like export factor 1 |
| NXT2 | nuclear transport factor 2-like export factor 2 |
| PPAN | peter pan homolog (Drosophila) |
| PA2G4 | proliferation-associated 2G4, 38kDa |
| PAK1IP1 | PAK1 interacting protein 1 |
| PAPD5 | PAP associated domain containing 5 |
| PARN | poly(A)-specific ribonuclease |
| PDCD11 | programmed cell death 11 |
| PELP1 | proline, glutamate and leucine rich protein 1 |
| PES1 | pescadillo ribosomal biogenesis factor 1 |
| PHAX | phosphorylated adaptor for RNA export |
| PHF6 | PHD finger protein 6 |
| PIH1D1 | PIH1 domain containing 1 |
| PINX1 | PIN2/TERF1 interacting, telomerase inhibitor 1 |
| PNO1 | partner of NOB1 homolog (S. cerevisiae) |
| PTRHD1 | peptidyl-tRNA hydrolase domain containing 1 |
| PWP1 | PWP1 homolog (S. cerevisiae) |
| PWP2 | PWP2 periodic tryptophan protein homolog (yeast) |
| RAI1 | retinoic acid induced 1 |
| RAN | RAN, member RAS oncogene family |
| RBM19 | RNA binding motif protein 19 |
| RBM28 | RNA binding motif protein 28 |
| RCL1 | RNA terminal phosphate cyclase-like 1 |
| REXO1 | REX1, RNA exonuclease 1 homolog (S. cerevisiae) |
| REXO2 | REX2, RNA exonuclease 2 homolog (S. cerevisiae) |
| REXO4 | REX4, RNA exonuclease 4 homolog (S. cerevisiae) |
| RIOK1 | RIO kinase 1 |
| RIOK2 | RIO kinase 2 |
| RNMTL1 | RNA methyltransferase like 1 |
| RPAP3 | RNA polymerase II associated protein 3 |
| RPF1 | ribosome production factor 1 homolog (S. cerevisiae) |
| RPF2 | ribosome production factor 2 homolog (S. cerevisiae) |
| RPS19BP1 | ribosomal protein S19 binding protein 1 |
| RRP1 | ribosomal RNA processing 1 homolog (S. cerevisiae) |
| RRP12 | ribosomal RNA processing 12 homolog (S. cerevisiae) |
| RRP36 | ribosomal RNA processing 36 homolog (S. cerevisiae) |
| RRP7A | ribosomal RNA processing 7 homolog A (S. cerevisiae) |
| RRP8 | ribosomal RNA processing 8, methyltransferase, homolog (yeast) |
| RRP9 | ribosomal RNA processing 9, small subunit (SSU) processome component, homolog (yeast) |
| RRS1 | RRS1 ribosome biogenesis regulator homolog (S. cerevisiae) |
| RSAD1 | radical S-adenosyl methionine domain containing 1 |
| RUVBL1 | RuvB-like 1 (E. coli) |
| RUVBL2 | RuvB-like 2 (E. coli) |
| SBDS | Shwachman-Bodian-Diamond syndrome |
| SDAD1 | SDA1 domain containing 1 |
| SHQ1 | SHQ1, H/ACA ribonucleoprotein assembly factor |
| SKIV2L2 | superkiller viralicidic activity 2-like 2 (S. cerevisiae) |
| SLFN14 | schlafen family member 14 |
| SPATA5 | spermatogenesis associated 5 |
| SPATA5L1 | spermatogenesis associated 5-like 1 |
| FTSJ3 | FtsJ homolog 3 (E. coli) |
| TAF9 | TAF9 RNA polymerase II, TATA box binding protein (TBP)-associated factor, 32kDa |
| TBL3 | transducin (beta)-like 3 |
| TCF25 | transcription factor 25 (basic helix-loop-helix) |
| TCOF1 | Treacher Collins-Franceschetti syndrome 1 |

| | |
|---|---|
| TEX10 | testis expressed 10 |
| TFB1M | transcription factor B1, mitochondrial |
| TGS1 | trimethylguanosine synthase 1 |
| TMA16 | translation machinery associated 16 homolog (S. cerevisiae) |
| TMED5 | transmembrane emp24 protein transport domain containing 5 |
| TNPO1 | transportin 1 |
| TSR1 | TSR1, 20S rRNA accumulation, homolog (S. cerevisiae) |
| TSR2 | TSR2, 20S rRNA accumulation, homolog (S. cerevisiae) |
| TSR3 | TSR3, 20S rRNA accumulation, homolog (S. cerevisiae) |
| TXNL4A | thioredoxin-like 4A |
| UFD1L | ubiquitin fusion degradation 1 like (yeast) |
| URB1 | URB1 ribosome biogenesis 1 homolog (S. cerevisiae) |
| URB2 | URB2 ribosome biogenesis 2 homolog (S. cerevisiae) |
| UTP11L | UTP11-like, U3 small nucleolar ribonucleoprotein, (yeast) |
| UTP14A | UTP14, U3 small nucleolar ribonucleoprotein, homolog A (yeast) |
| UTP15 | UTP15, U3 small nucleolar ribonucleoprotein, homolog (S. cerevisiae) |
| UTP18 | UTP18 small subunit (SSU) processome component homolog (yeast) |
| UTP20 | UTP20, small subunit (SSU) processome component, homolog (yeast) |
| UTP23 | UTP23, small subunit (SSU) processome component, homolog (yeast) |
| UTP3 | UTP3, small subunit (SSU) processome component, homolog (S. cerevisiae) |
| UTP6 | UTP6, small subunit (SSU) processome component, homolog (yeast) |
| VCP | valosin containing protein |
| WDR12 | WD repeat domain 12 |
| WDR18 | WD repeat domain 18 |
| WDR36 | WD repeat domain 36 |
| WDR43 | WD repeat domain 43 |
| WDR46 | WD repeat domain 46 |
| WDR74 | WD repeat domain 74 |
| WDR75 | WD repeat domain 75 |
| WRAP53 | WD repeat containing, antisense to TP53 |
| XPO1 | exportin 1 (CRM1 homolog, yeast) |
| XRN1 | 5'-3' exoribonuclease 1 |
| XRN2 | 5'-3' exoribonuclease 2 |
| ZCCHC4 | zinc finger, CCHC domain containing 4 |
| ZNHIT3 | zinc finger, HIT-type containing 3 |
| ZNHIT6 | zinc finger, HIT-type containing 6 |
| BMT2 | base methyltransferase of 25S rRNA 2 homolog |
| BUD23 | BUD23 rRNA methyltransferase and ribosome maturation factor |
| CCDC86 | coiled-coil domain containing 86 |
| DHX8 | DEAH (Asp-Glu-Ala-His) box polypeptide 8 |
| NOP53 | glioma tumor suppressor candidate region gene 2 |
| NOPCHAP1 | chromosome 12 open reading frame 45 |
| NLE1 | notchless homolog 1 (Drosophila) |
| SLX9 | family with sequence similarity 207, member A |
| NOL11 | nucleolar protein 11 |
| AQR | aquarius homolog (mouse) |
| C1D | C1D nuclear receptor corepressor |
| RSL1D1 | ribosomal L1 domain containing 1 |
| NIFK | MKI67 (FHA domain) interacting nucleolar phosphoprotein |
| ZNF622 | zinc finger protein 622 |
| PDCD2 | programmed cell death 2 |
| PDCD2L | programmed cell death 2-like |
| GEMIN5 | gem (nuclear organelle) associated protein 5 |
| RBM34 | AT rich interactive domain 4B (RBP1-like) // RNA binding motif protein 34 |
| BOP1 | BOP1 ribosomal biogenesis factor |

**Supplementary table S2.**

Characteristics of PAM50 and intrinsic breast cancer subtype cohorts from TCGA dataset

| | **PAM50 (n = 739)** | **Intrinsic breast cancer subtype (n = 712)** |
|---|---|---|
| **Age (years)** | | |
| Mean (SD) | 58.1 (13.2) | 58.1 (13.3) |
| Median [Min: Max] | 58.0 [26.0 : 90.0] | 58.0 [26.0 : 90.0] |
| **Gender** | | |
| Female | 739 (100%) | 712 (100%) |
| Male | 0 (0%) | 0 (0%) |
| **Tumour size** | | |
| T1 | 195 (26.4%) | 186 (26.1%) |
| T2 | 408 (55.2%) | 421 (59.1%) |
| T3 | 71 (9.6%) | 77 (10.8%) |
| T4 | 24 (3.2%) | 25 (3.5%) |
| TX | 2 (0.3%) | 3 (0.4%) |
| Missing | 39 (5.3%) | 0 (0%) |
| **Node invasion** | | |
| N0 | 345 (46.7%) | 354 (49.7%) |
| N1 | 233 (31.5%) | 235 (33.0%) |
| Missing | 161 (21.8%) | 123 (17.3%) |
| **Metastasis** | | |
| M0 | 684 (92.6%) | 695 (97.6%) |
| M1 | 12 (1.6%) | 12 (1.7%) |
| Missing | 43 (5.8%) | 5 (0.7%) |
| **Stage** | | |
| Stage I | 112 (15.2%) | 109 (15.3%) |
| Stage II | 299 (40.5%) | 305 (42.8%) |
| Stage III | 103 (13.9%) | 100 (14.0%) |
| Stage IV | 4 (0.5%) | 4 (0.6%) |
| Missing | 221 (29.9%) | 194 (27.2%) |
| **Histological type** | | |
| Infiltrating Ductal Carcinoma | 611 (82.7%) | 580 (81.5%) |

| | | |
|---|---|---|
| Infiltrating Lobular Carcinoma | 60 (8.1%) | 68 (9.6%) |
| Medullary Carcinoma | 4 (0.5%) | 3 (0.4%) |
| Mixed Histology | 28 (3.8%) | 23 (3.2%) |
| Mucinous Carcinoma | 5 (0.7%) | 5 (0.7%) |
| Other | 30 (4.1%) | 32 (4.5%) |
| Missing | 1 (0.1%) | 1 (0.1%) |
| **Neoadjuvant treatment** | | |
| Yes | 0 (0%) | 0 (0%) |
| No | 739 (100%) | 712 (100%) |

**Supplementary Table S3.**
Comparison of TCGA breast cancer sample classifications

| Intrinsic status#/PAM50$ | Luminal A (n = 362) | Luminal B (n = 172) | HER2 (n = 68) | TNBC (n = 137) | NA (n = 189) |
|---|---|---|---|---|---|
| **Luminal A (n=486)** | **289** | 123 | 5 | 19 | 50 |
| **Luminal B (n=74)** | 21 | **27** | 19 | 1 | 6 |
| **HER2 (n=30)** | 0 | 1 | **26** | 2 | 1 |
| **TNBC (n=122)** | 3 | 1 | 10 | **101** | 7 |
| **NA (=216)** | 49 | 20 | 8 | 14 | **125** |

$: PAM50 classification relying on transcriptomics is directly given by the TCGA database
#: Intrinsic breast cancer status classification relying on detection of oestrogen, progesterone and HER2 receptors was determined using the positive or negative IHC status given by the TCGA database

**Supplementary table S4.**

Primary antibodies used in Western blot

| Protein | M.W. | Dilution | Provider / reference |
|---|---|---|---|
| PARP | 120 | 1/1000 | Cell Signalling - 9532S |
| PARP 1 total | 133 | 1/1000 | Abcam - ab137653 |
| Cleaved PARP | 89 | 1/1000 | Cell signalling - 5625 |
| Caspase 3 | 35 | 1/100 | Enzo - ALX-804-305-C100 |
| Cleaved caspase 3 | 17 | 1/1000 | Cell signalling - 9661 |
| Fibrillarin | 37 | 1/1000 | Abcam - Ab166630 |
| Ku80 | 80 | 1/2000 | Abcam - Ab119935 |
| β-Actin | 42 | 1/5000 | Sigma - A5441 |
| H3 | 17 | 1/5000 | Abcam - ab1791 |
| γH2A.X | 17 | 1/1000 | Millipore - 05-636 |
| Phospho-Histone H3 (ser10) | 17 | 1/1000 | Cell Signalling - 9701 |

**Supplementary table S5.**

Antibodies used for immunofluorescence

| Protein | Dilution | Provider / reference |
|---|---|---|
| FBL | 1/4000 | Abcam - Ab5821 |
| Phospho-H3(S10) | 1/1000 | Cell Signalling - 9701S |
| | 1/1000 | Merck - 06-570 |
| Gamma H2Ax | 1/6000 | Merck - 05-636 |
| 53BP1 | 1/500 | Cell Signalling - 4937-S |
| Alexa-Fluor 488 anti-Rabbit | 1/1000 | Invitrogen - A11008 |
| Alexa-Fluor 488 anti-Mouse | 1/1000 | Invitrogen - A11001 |
| Alexa-Fluor 555 anti-Rabbit | 1/1000 | Invitrogen - A21428 |
| Alexa-Fluor 555 anti-Mouse | 1/1000 | Invitrogen - A21424 |
| Alexa-Fluor 647 anti-Rabbit | 1/1000 | Invitrogen - A21245 |
| Alexa-Fluor 647 anti-Mouse | 1/1000 | Invitrogen - A21235 |
